# Designing epitope-focused vaccines *via* antigen reorientation

**DOI:** 10.1101/2022.12.20.521291

**Authors:** Duo Xu, Chunfeng Li, Ashley Utz, Payton A.B. Weidenbacher, Shaogeng Tang, Mrinmoy Sanyal, Bali Pulendran, Peter S. Kim

**Affiliations:** Department of Biochemistry, Stanford University School of Medicine, Stanford, CA, USA; Sarafan ChEM-H, Stanford University, Stanford, CA, USA; Institute for Immunity, Transplantation and Infection, Stanford University, Stanford, CA, USA; Stanford Medical Scientist Training Program, Stanford University School of Medicine, Stanford, CA, USA; Stanford Biophysics Program, Stanford University School of Medicine, Stanford, CA, USA; Department of Chemistry, Stanford University, Stanford, CA, USA; Department of Pathology, Stanford University School of Medicine, Stanford, CA, USA; Department of Microbiology and Immunology, Stanford University School of Medicine, Stanford, CA, USA; Chan Zuckerberg Biohub, San Francisco, CA, USA

## Abstract

A major challenge in vaccine development, especially against rapidly evolving viruses, is the ability to focus the immune response toward evolutionarily conserved antigenic regions to confer broad protection. For example, while many broadly neutralizing antibodies against influenza have been found to target the highly conserved stem region of hemagglutinin (HA-stem), the immune response to seasonal influenza vaccines is predominantly directed to the immunodominant but variable head region (HA-head), leading to narrow-spectrum efficacy. Here, we first introduce an approach to controlling antigen orientation based on the site-specific insertion of short stretches of aspartate residues (oligoD) that facilitates antigen-binding to alum adjuvants. We demonstrate the generalizability of this approach to antigens from the Ebola virus, SARS-CoV-2, and influenza and observe enhanced antibody responses following immunization in all cases. Next, we use this approach to reorient HA in an “upside down” configuration, which we envision increases HA-stem exposure, therefore also improving its immunogenicity compared to HA-head. When applied to HA of H2N2 A/Japan/305/1957, the reoriented H2 HA (reoH2HA) on alum induced a stem-directed antibody response that cross-reacted with both group 1 and 2 influenza A HAs. Our results demonstrate the possibility and benefits of antigen reorientation *via* oligoD insertion, which represents a generalizable immunofocusing approach readily applicable for designing epitope-focused vaccine candidates.

**GRAPHICAL ABSTRACT:** 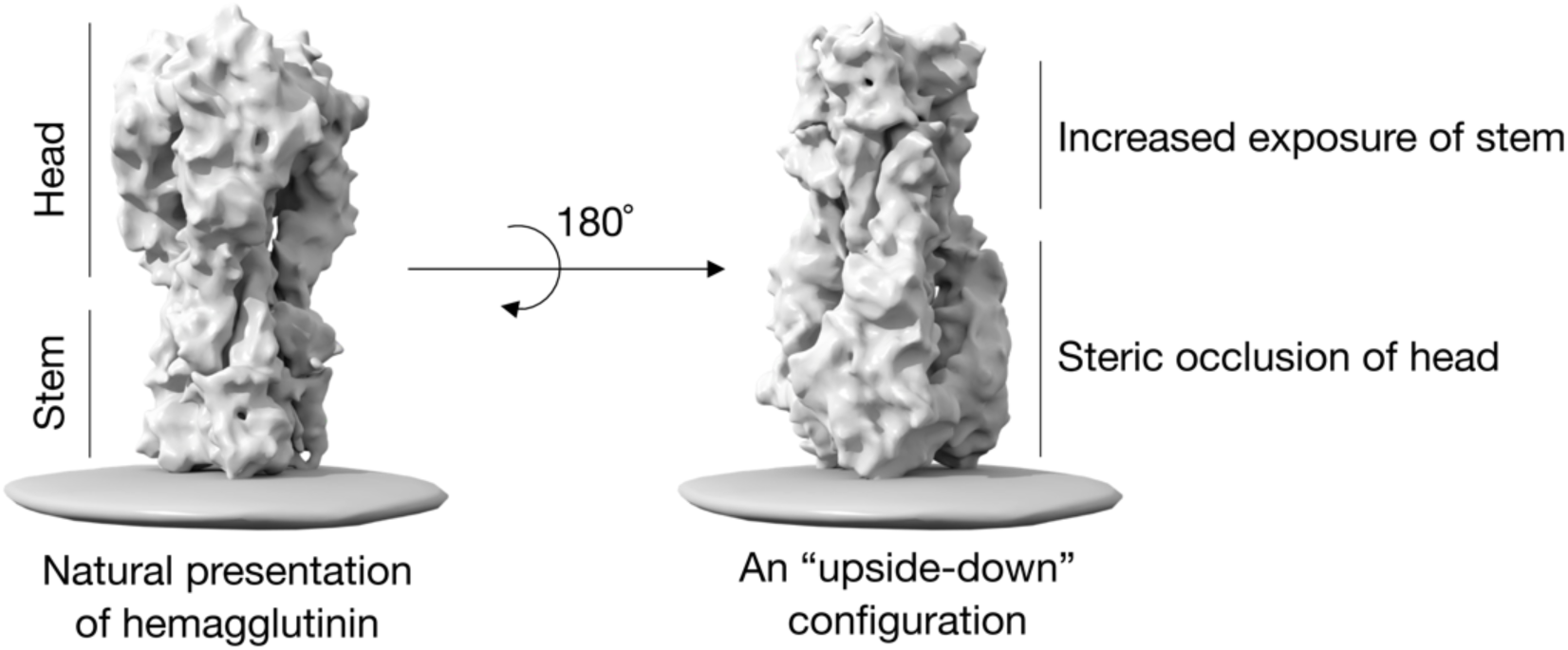

Seasonal influenza vaccines induce a biased antibody response against the variable head of hemagglutinin, whereas conserved epitopes on the stem are a target for universal vaccines. Here we show that reorienting HA in an “upside-down” configuration sterically occludes the head and redirects the antibody response to the more exposed stem, thereby inducing broad cross-reactivity against hemagglutinins from diverse influenza strains.

## INTRODUCTION

The rapid evolution of viruses like influenza and SARS-CoV-2 leads to periodic outbreaks with varying severity and represents a serious public health concern^1–3^. In the case of influenza, current vaccines elicit strain-specific antibody responses against the immunodominant but variable head region of HA (HA-head) that undergoes antigenic drift, necessitating annual updates of seasonal influenza vaccines^4,5^. When a novel zoonotic influenza virus acquires the ability of sustained human-to-human transmission, existing vaccine-induced immunity is unlikely to confer protection^6^. Such risks highlight the importance of pandemic preparedness and the urgent need to generate a universal vaccine that protects against diverse influenza strains^7^.

A potential strategy for developing more broadly protective vaccines is to focus the immune response (referred to as immunofocusing) on antigenic regions that are less likely to evolve. Since the stem region of HA (HA-stem) contains highly conserved epitopes across different influenza subtypes^8,9^, focusing the antibody response on HA-stem and away from the immunodominant HA-head is a promising approach. Human monoclonal antibodies (mAbs) targeting HA-stem show broad neutralizing activity and protect animals from lethal challenges^10–14^. HA-stem-directed antibodies also correlate with protection from infection and disease in humans^15^.

Current immunofocusing approaches include sequential immunization^16–19^ or mosaic display^20,21^ of heterologous HAs, or chimeric HAs that contain the same HA-stem but varying HA-head to stimulate a stem-directed response^22^. These approaches require complicated vaccination regimens and do not directly address the immunodominance of HA-head^23^. Other immunofocusing approaches include masking HA-head with glycosylation^24,25^ or PEGylation^26^, or structure-based design of HA-stem-only immunogens^27–30^. While epitope masking often leads to reduced immunogenicity *in vivo*^25,26^, the generation of HA-stem-only proteins requires laborious rounds of design, screening, and optimization of antigens to retain the proper presentation of conformational epitopes. Ultimately, although these strategies have found some success in inducing cross-reactivity among a particular phylogenetic group (i.e., group 1 or 2) of influenza A HAs, it remains a challenge to elicit a consistent response to both groups from a single immunogen.

Here, we investigate the hypothesis that antigen reorientation is a straightforward and effective immunofocusing strategy. More specifically, we test if the reorientation of HA in an “upside-down” configuration from its natural presentation on influenza virions would sterically occlude HA-head and redirect the antibody response to the more exposed HA-stem.

Interestingly, stem-directed (but not head-directed) mAbs isolated from vaccinated individuals exhibit significantly reduced binding to the whole virus as compared to recombinant HA proteins^31^. We hypothesize that steric occlusion leads to poor accessibility of HA-stem^32^, thereby biasing antibody responses toward HA-head during natural infection^31^.

To control antigen orientation, we first develop an approach based on the site-specific insertion of short stretches of aspartate residues (oligoD) into different protein antigens to facilitate their binding to alum, a well-established vaccine adjuvant that has been used for over 90 years^33,34^. The surface of alum allows the presentation of oligoD-modified antigens in a pre-defined manner. While methods to afford increased antigen-binding to alum have been developed previously, including modifications with terminal phosphate^35^ or phosphonate groups^36^ or phosphoserine peptides^37,38^, these approaches usually require the synthesis of alum-binding molecules followed by chemical coupling and post-modification purification procedures. In contrast, oligoD insertion is a simple, one-step process pre-programmed by molecular cloning.

To validate our proposed approach, we inserted oligoD into the C-terminus of Ebola glycoprotein and showed that oligoD insertion increased antigen-binding to alum and elicited robust germinal center and neutralizing antibody responses. To demonstrate the simplicity and generalizability of our approach, we also inserted oligoD into two other protein antigens (SARS-CoV-2 spike and HA of A/New Caledonia/20/99) and found that oligoD-modified antigens induced significantly stronger neutralizing antibody responses compared to wild-type antigens.

After establishing that oligoD insertion mediated antigen-binding to alum, we identified a permissive site on the head of an H2 HA (A/Japan/305/1957) for oligoD insertion. We choose this H2 HA because H2 HAs contain a bulky phenylalanine residue on its stem region that expands the breadth of B cell responses upon vaccination compared to HAs from other subtypes^39^. We showed that oligoD insertion into H2 HA resulted in the trivalent anchoring of its head to alum at specific locations on each HA trimer in an “upside down” configuration. Immunization with this reoriented H2 HA (reoH2HA) on alum elicited a stem-directed and cross-reactive antibody response to diverse HA subtypes from both group 1 and 2. Our results demonstrate the potential of designing epitope-focused vaccine candidates *via* antigen reorientation.

## RESULTS

### OligoD insertion enables antigen-binding to alum and enhances neutralizing antibody responses

To determine the length of oligoD required for alum-binding, we first used the ectodomain of Ebola glycoprotein (GP) as a test case and genetically inserted 2, 4, 8 or 12 repeating units of aspartates (abbreviated as 2D, 4D, 8D or 12D, respectively) at its C-terminus (**Fig. 1a**). OligoD-modified GP proteins were expressed in Expi293F cells *via* transient transfection and purified to homogeneity (**Fig. S1a**). OligoD insertion did not change the thermal melting profile (**Fig. 1b**) or melting temperature (Tm, 58°C, **Fig. S1b**) of GP, suggesting a negligible perturbation to its native structure. In addition, we selected a panel of five GP-specific monoclonal antibodies (mAb114, c13C6, ADI-15742, KZ52 and ADI-16061) that recognize different epitopes on GP and examined their binding to wild-type or oligoD-modified GP using bio-layer interferometry (BLI, **Fig. S1c**). We found no noticeable differences in the binding profiles of these mAbs to all GP proteins (**Fig. 1c**), suggesting proper presentation of those conformational epitopes. Next, we measured the effect of oligoD insertions on GP-binding to alum (Alhydrogel^®^). We pre-mixed wild-type or oligoD-modified GP with alum (protein: alum, 1: 10, *w/w*) for one hour and then incubated the mixtures in the presence of naïve mouse serum (10%, *v/v*) at 37 °C for 24 hours (**Fig. S1d**). After incubation, alum pellets were rinsed extensively and subjected to protein gel electrophoresis, followed by Western-blot analysis to determine the amount of alum-bound GP. Whereas only a small fraction of wild-type GP was detected on the blot, an increasing amount of GP bound to alum as the number of aspartates increased from two to 12 (**Fig. 1d**). To quantify the amount of alum-bound GP, we collected the supernatant from GP-alum mixtures and measured the concentration of unbound GP by enzyme-linked immunosorbent assays (ELISAs, **Fig. S1e**). Consistent with the Western blot, higher GP-binding to alum correlated with increasing oligoD length (**Fig. 1e**, **Fig. S1f**). The presence of naïve mouse serum did not affect binding of GP-8D or GP-12D to alum. This interaction between oligoD and alum also did not affect the thermal stability of GP proteins (**Fig. S1g,h**). Besides the C-terminus, we identified three flexible loop regions (after residue R200, T294 or A309) on GP that readily accommodated oligoD insertion (**Fig. S2a,b**). Thermal melting profiles and T_m_ of the resulting oligoD-modified GP proteins were identical to those of wild-type GP (**Fig. S2c**). While both GP-8D and GP-12D exhibited ∼100% binding to alum, only 12D insertion afforded ∼100% GP-binding to alum when inserted into each of these flexible loop regions (**Fig. S2d**). Therefore, we inserted 12D into antigens to promote maximal binding to alum for subsequent immunization studies.

**Figure 1.**
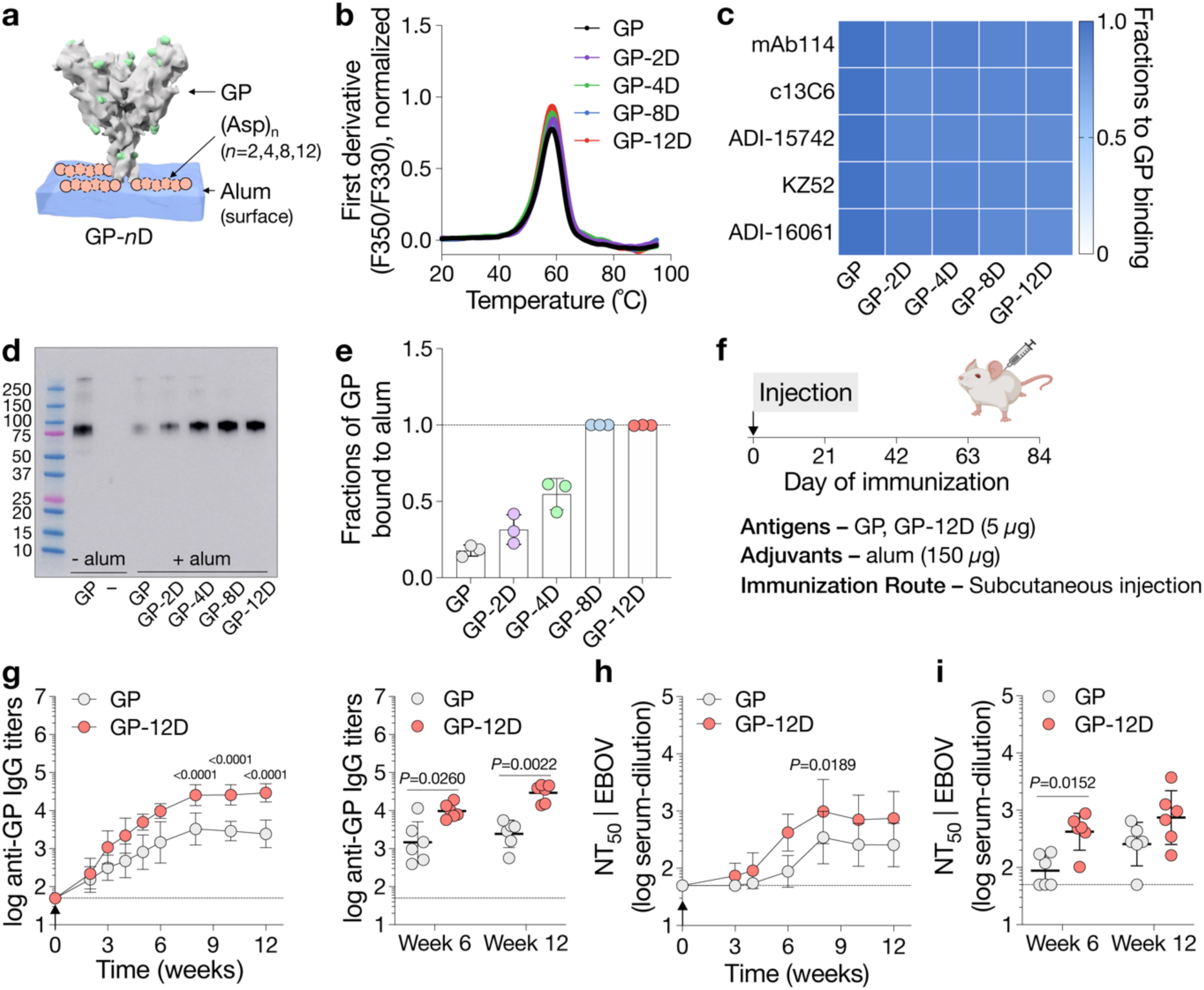
OligoD insertion enables antigen-binding to alum and enhances neutralizing antibody responses. **a**, Site-specific insertion of oligoD of different lengths (2, 4, 8 or 12 aspartate residues, pink) at the C-terminus of Ebola GP (PDB ID: 5JQ3). **b**, Thermal melting profiles of wild-type or oligoD-modified GP measured by differential scanning fluorimetry. **c**, Analysis of GP-specific mAb binding of wild-type or oligoD-modified GP by bio-layer interferometry (BLI). Shifts in nanometers of mAb-binding to oligoD-modified GP were normalized as a fraction of the shifts to wild-type GP binding. Fractions were calculated based on data in **Fig. S1c. d,e**, Detection and quantification of alum-bound GP by Western-blot analysis (**d**) and ELISA (**e**). Wild-type or oligoD-modified GP proteins were pre-mixed with alum and then incubated in PBS containing naïve mouse serum for 24 hour at 37 °C. Upon separation, alum-bound GP was detected by mAb114 on the Western blot, while unbound GP was quantified by ELISA as in **Fig. S1f**. Dashed line indicates 100% binding to alum in **e**. Data are presented as mean ± s.d. (*n*=3 samples per group). **f**, A single-dose immunization study with wild-type GP or GP-12D adjuvanted with alum *via* subcutaneous injection in BALB/c mice (*n*=6 mice per group). **g**, Serum GP-specific IgG titers over time. Antibody titers at week 6 and 12 post-immunization are plotted on the right for comparison of the two groups. **h**, Serum neutralization titers (NT_50_) against Ebola GP-pseudotyped lentiviruses (EBOV) over time. **i**, NT_50_ of week 6 and 12 from the two groups were shown for comparison. Dashed lines indicate the limit of quantification in **g-i**. Data are presented as geometric mean ± s.d. of the log-transformed values (**g-i**). Comparison of IgG titers (**g**) or NT_50_ (**h**) over time was performed using two-way ANOVA followed by a Bonferroni test. Comparison of two groups was performed using the two-tailed Mann–Whitney U test (**g,h**). *P* values of 0.05 or less were considered significant and plotted.

To evaluate the immunogenicity of oligoD-modified GP, we immunized mice with either wild-type GP or GP-12D adjuvanted with alum *via* subcutaneous injection (**Fig. 1f**). A single dose of GP with alum elicited a relatively weak GP-specific antibody response, while GP-12D with alum stimulated a much stronger response with almost tenfold higher IgG titers six weeks post-immunization (**Fig. 1g**). At six and eight weeks post-immunization, we detected very low IgG titers against oligoD or other C-terminal affinity tags that were used for purification purposes (**Fig. S3a**). Next, we generated Ebola GP-pseudotyped lentiviruses (EBOV, **Fig. S3b**) to analyze the neutralizing antibody response. Consistent with the IgG titers, GP-12D induced a three-to five-fold stronger neutralizing antibody response than GP did (**Fig. 1h**). While all mice immunized with GP-12D adjuvanted with alum developed neutralizing responses at week six (**Fig. S3c**), only half of the mice immunized with GP did, though the titers were lower. Even at week 12, one mouse immunized with GP did not show any neutralizing activity (**Fig. 1i**, **Fig. S3d**). Taken together, oligoD insertion increased antigen-binding to alum and significantly enhanced neutralizing antibody responses in animal immunizations compared to wild-type GP.

### OligoD-modified antigens stimulate robust germinal center responses

To understand the mechanism underlying the enhanced antibody response elicited by GP-12D, we immunized mice with wild-type GP or GP-12D and analyzed the germinal center (GC) responses in draining lymph nodes at 7, 14 or 21 days post-immunization using flow cytometry^40^ (**Fig. 2a**). As with the previous immunization (**Fig. 1g**), a single injection of GP-12D with alum elicited a stronger antibody response than GP with alum did (**Fig. 2b,c**). Use of alum also led to a Th2-biased response (IgG1) in both groups with negligible titers of IgG2a or IgG2b (**Fig. S4a**).

**Figure 2.**
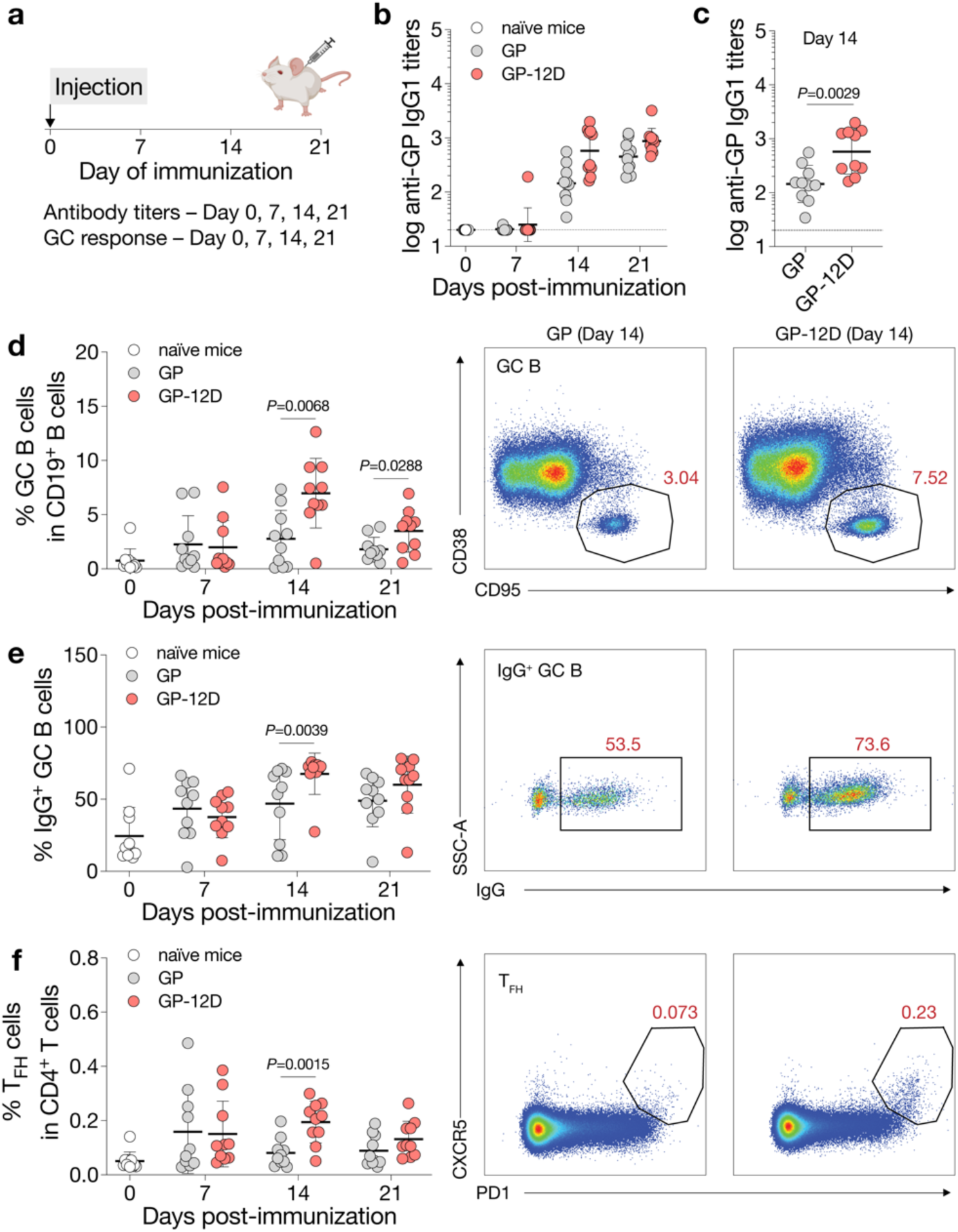
OligoD-modified GP stimulates a robust germinal center response. **a**, Immunization schedule for the analysis of GC responses in draining lymph nodes. BALB/c mice were immunized with wild-type GP or GP-12D adjuvanted with alum (*n*=10 mice per antigen per time point). Antibody titers and GC responses were measured pre-immunization and 7, 14 or 21 days post-immunization. **b**, Serum GP-specific IgG1 titers over time. **c**, IgG1 titers of day 14 were shown for comparison. Dashed lines indicate the limit of quantification in **b,c**. **d-f**, Analysis of GC B cell (**d**), IgG^+^ GC B cell (**e**) and T_FH_ cell (**f**) responses after immunization. Data are presented as geometric mean ± s.d. of the log-transformed values (**b,c**) or mean ± s.d. (**d-f**). Comparison of two groups was performed using the two-tailed Mann–Whitney U test (**c-f**). *P* values of 0.05 or less were considered significant and plotted.

Besides the serological response, we examined the responses of GC B cells (CD19^+^CD95^+^CD38^−^), IgG^+^ GC B cells and T follicular helper cells (T_FH_, CD3^+^CD4^+^PD1^+^CXCR5^+^) in the draining lymph nodes (**Fig. S4b**). Compared to GP with alum, GP-12D with alum induced an extended GC response with a substantially higher magnitude of GC B (**Fig. 2d**), IgG^+^ GC B (**Fig. 2e**) and T_FH_ cells (**Fig. 2f**), which all peaked at 14 days post-immunization. These data suggest that the higher antibody titers elicited by GP-12D with alum resulted from stimulation of a robust GC response.

### OligoD insertion is generalizable to other protein antigens to enhance antibody responses

Given the simplicity of our approach, we applied oligoD insertion to two other protein antigens for additional validation. We first inserted oligoD into the C-terminus of SARS-CoV-2 spike to generate spike-12D (**Fig. 3a**). Wild-type spike and spike-12D were transiently expressed and purified to homogeneity (**Fig. S5a**). In both the absence (**Fig. S5b**) and presence of alum (**Fig. S5c**), spike-12D shared the same thermal melting profile and T_m_ (44°C and 52°C) with wild-type spike. We also found nearly identical binding profiles for ACE2-Fc and three spike-specific mAbs (COVA2-15, CB6 and CR3022) against wild-type spike or spike-12D (**Fig. 3b**, **Fig. S5d**). Next, we immunized mice with wild-type spike or spike-12D adjuvanted with alum *via* subcutaneous injection (**Fig. 3c**). A three-dose regimen of spike-12D with alum elicited a significantly higher antibody response than wild-type spike with alum did against both spike (**Fig. 3d**) and its receptor-binding domain (RBD, **Fig. 3e**). Immunization of spike-12D with alum also greatly enhanced the neutralizing response against spike-pseudotyped lentiviruses (**Fig. 3f**, **Fig. S6a,b**). Despite the less than twofold difference in IgG titers at week nine, the neutralization titers (NT_50_) of the spike-12D group were over fivefold higher than those of the spike group. When we further assessed the neutralization of other SARS-CoV-2 variants of concern (**Fig. S6c-g**), we found consistently higher neutralizing activity from antisera elicited by spike-12D (**Fig. 3g**).

**Figure 3.**
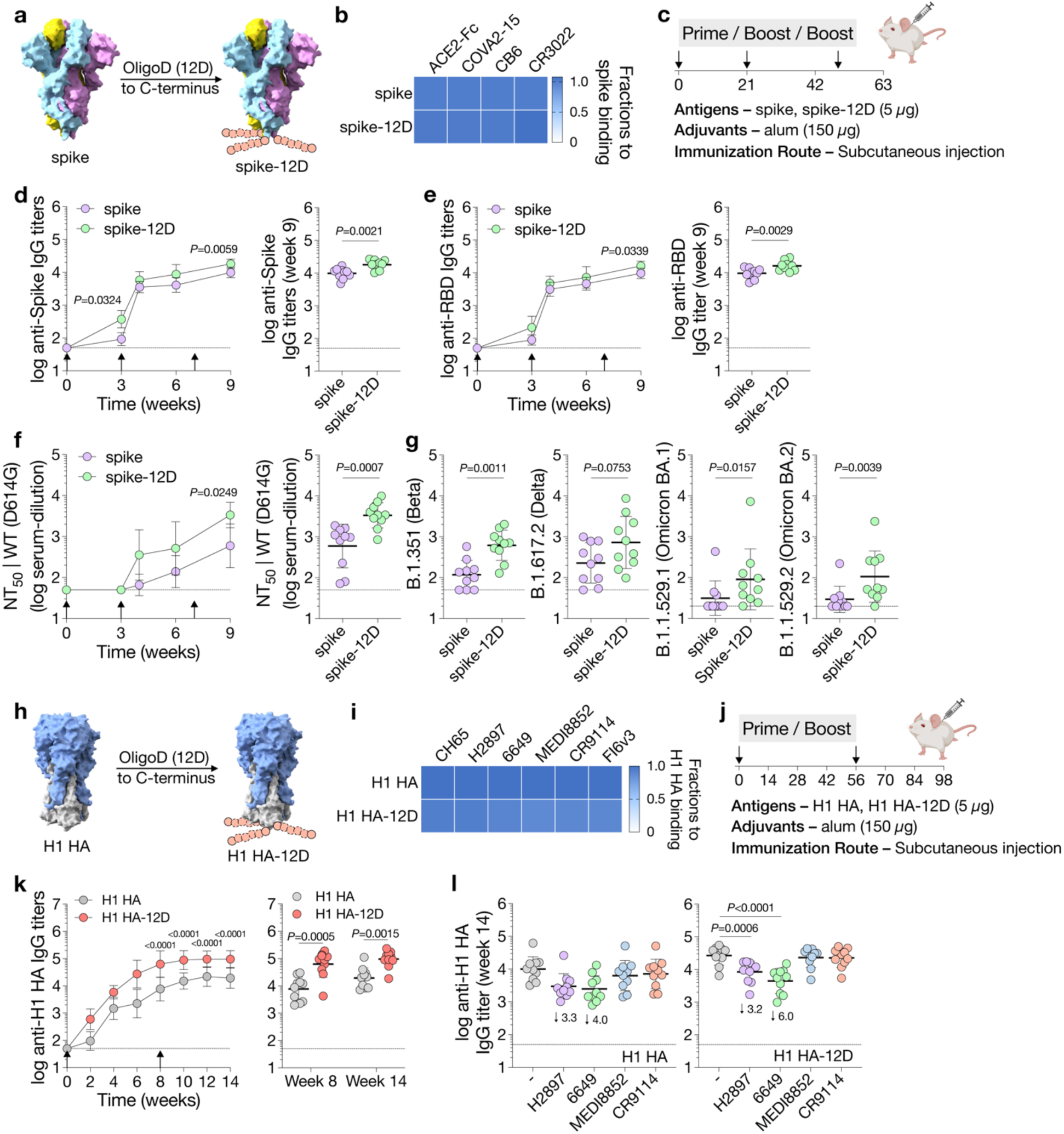
OligoD insertion is generalizable to other protein antigens. **a**, Insertion of oligoD (12D) into the C-terminus of SARS-CoV-2 spike (PDB ID: 6VXX). **b**, BLI binding analysis of wild-type spike or spike-12D with ACE2-Fc and three spike-specific mAbs. **c**, A three-dose immunization study with wild-type spike or spike-12D adjuvanted with alum *via* subcutaneous injection in BALB/c mice (*n*=10 mice per group). **d,e**, Serum spike-(**d**) or RBD-specific (**e**) IgG titers over time. Endpoint (week 9) antibody titers are plotted on the right for comparison of the two groups. **f**, Serum NT_50_ against SARS-CoV-2 spike-pseudotyped lentiviruses over time. Endpoint (week 9) NT_50_ are plotted on the right for comparison of the two groups. **g**, Endpoint (week 9) NT_50_ against SARS-CoV-2 variants of concern from the two groups. **h**, Insertion of oligoD (12D) into the C-terminus of H1 HA (PDB ID: 1RU7 for reference). **i**, HA-specific mAb binding analysis of wild-type or oligoD-modified H1 HA by BLI. **j**, A prime-boost immunization study with wild-type H1 HA or H1 HA-12D adjuvanted with alum *via* subcutaneous injection in BALB/c mice (*n*=10 mice per group). **k**, Serum H1 HA-specific IgG titers over time. Antibody titers of week 8 and week 14 from the two groups are plotted on the right for comparison. **l**, Serum binding titers to H1 HA in the presence of competing mAbs. Fold-change is indicated by arrows with numbers. Dashed lines indicate the limit of quantification in **d-g,k,l**. Arrows in **d-f** and **k** indicate prime and boost immunizations. Data are presented as geometric mean ± s.d. of the log-transformed values. Comparison of IgG titers (**d,e,k**) or NT_50_ (**f**) over time was performed using two-way ANOVA followed by a Bonferroni test. Comparison of two groups was performed using the two-tailed Mann–Whitney U test (**d-g,k,l**). Comparison of multiple groups was performed using one-way ANOVA followed by a Bonferroni test (**l**). *P* values of 0.05 or less were considered significant and plotted.

Besides SARS-CoV-2 spike, we inserted oligoD into the C-terminus of an H1 HA (A/New Caledonia/20/99) (**Fig. 3h**). Similar to the modifications in GP and spike, oligoD insertion did not affect the expression (**Fig. S7a**) or thermal stability (**Fig. S7b**) of H1 HA. The binding profiles with different HA-specific mAbs (CH65, H2897, 6649, MEDI8852, CR9114 and FI6v3), as determined by BLI, also suggested the proper presentation of conformational epitopes on H1 HA-12D (**Fig. S7c, Fig. 3i**). A prime-boost immunization of H1 HA-12D with alum elicited substantially higher (five-to tenfold) HA-specific IgG titers than H1 HA with alum did (**Fig. 3j,k**). To determine whether the antisera targeted HA-head or HA-stem, we performed competition ELISA, in which antigen-coated ELISA plates were pre-incubated with a saturating concentration of individual HA-specific mAbs before incubation with antisera. We observed a clear competition between antisera and HA-head-directed mAbs (H2897 or 6649) with a more pronounced change in the H1 HA-12D group (**Fig. 3l**). In contrast, there was negligible competition with HA-stem-directed antibodies (MEDI8852 or CR9114), suggesting that oligoD insertion into the C-terminus of HA only boosted antibody responses toward the immunodominant HA-head. Nonetheless, higher antibody titers elicited by H1 HA-12D correlated with an increased breadth against two other group 1 HAs, namely H2 (A/Japan/305/1957) and H5 (A/Vietnam/1203/2004) HAs, presumably toward conserved epitopes on HA-head (**Fig. S7d**).

### OligoD insertion into the head of H2 HA reorients it on alum to afford immunofocusing

Having validated using three protein antigens that oligoD insertion enabled their binding to alum, we then attempted to insert oligoD into HA-head to anchor this region on alum and redirect antibody responses to the newly exposed HA-stem (**Fig. 4a**). We chose to insert oligoD into an H2 HA because the bulky phenylalanine (F45_HA2_) in its HA-stem expands the cross-reactivity of stem-binding B cells^39^. Although oligoD insertions into HA-head often led to substantially reduced protein expression levels (**Fig. S8a**), insertion of 12D after residue S156 in the head of H2 HA (A/Japan/305/1957) was successful and produced reoH2HA (**Fig. S8b**).

**Figure 4.**
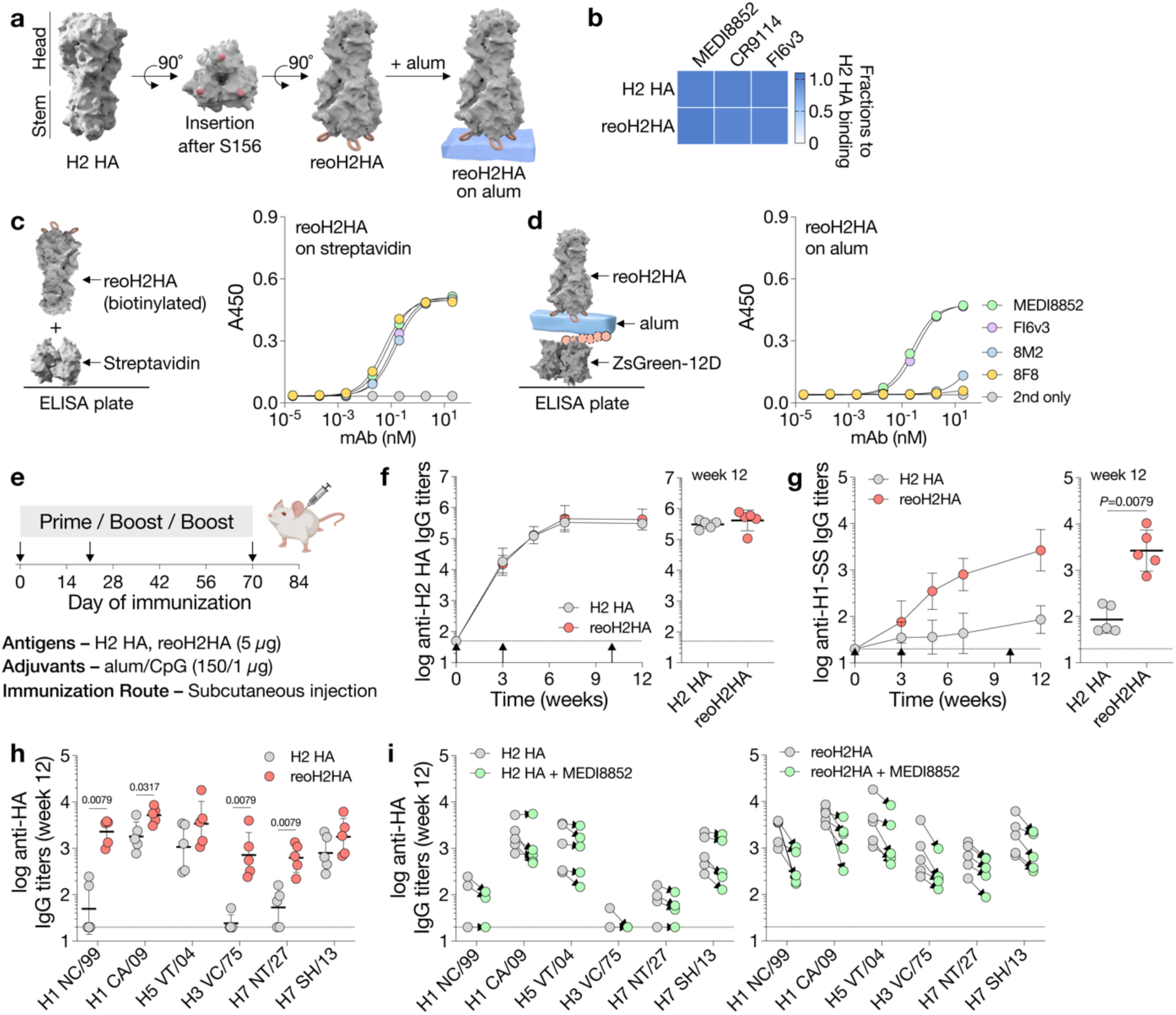
Reorientation of H2 HA enabled immunofocusing on HA-stem. **a**, Insertion of oligoD (12D) into the head of H2 HA (A/Japan/305/1957) after residue S156 (PDB ID: 2WRE). Three oligoD motifs on HA-head allowed for its trivalent anchoring on alum. **b**, HA-stem-specific mAb binding analysis of wild-type H2 HA or reoH2HA by BLI. **c**, Binding of stem-directed (MEDI8852 and FI6v3) or head-directed mAbs (8F8 and 8M2) to reoH2HA on streptavidin-(**c**) or alum-based (**d**) ELISA plates. (*n*=2 biological replicates of technical duplicates). **e**, A three-dose immunization study with wild-type H2 HA or reoH2HA adjuvanted with alum/CpG *via* subcutaneous injection in BALB/c mice (*n*=5 mice per group). **f,g**, Serum H2 HA-specific (**f**) or stem-specific (**g**) IgG titers over time. Stem-specific IgG titers were measured with the H1-stablized stem (H1-SS) protein. Endpoint (week 12) antibody titers from the two groups are plotted on the right for comparison. Arrows indicate prime and boost immunizations. **h**, Cross-reactive binding of group 1 (H1 NC/99, H1 CA/09, H2 JP/57, H5 VT/04) and group 2 HAs (H3 VC/75, H7 NT/27, H7 SH/13 – A/Shanghai/2/2013) by week-12 antisera. **i**, Serum binding titers to group 1 and 2 HAs in the presence of competing mAb – MEDI8852. Dashed lines indicate the limit of quantification in **f-i**. Data are presented as mean ± s.d. (**c,d**) or geometric mean ± s.d. of the log-transformed values (**f-i**). Comparison of two groups was performed using the two-tailed Mann–Whitney U test. *P* values of 0.05 or less were considered significant and plotted.

ReoH2HA maintained the thermal stability of its wild-type counterpart (**Fig. S8c**), including in the presence of alum (**Fig. S8d**). The insertion into the head of H2 HA also did not appreciably affect the binding profiles (**Fig. S8e, Fig. 4b**) or affinities (**Fig. S8f-h**) of stem-directed mAbs against the protein. To validate the “upside-down” orientation of oligoD-modified HA on alum, we measured binding of head-and stem-directed mAbs to reoH2HA on streptavidin-or alum-coated ELISA plates (**Fig. 4c,d**). Both head-(8F8 and 8M2^41^) and stem-directed mAbs (MEDI8852 and FI6v3) bound to reoH2HA with high affinity when C-terminal biotinylated reoH2HA was adsorbed on streptavidin-coated ELISA plates (**Fig. 4c**). In contrast, only stem-directed mAbs bound to reoH2HA when it was adsorbed on alum (**Fig. 4d**). Loss of binding of head-directed mAbs suggests that reoH2HA adopted an “upside down” configuration on alum, where epitopes on HA-head were no longer accessible. As a control, when we tested mAb binding to H2 HA with 12D inserted into its C-terminus, binding profiles were almost identical between streptavidin-and alum-coated plates (**Fig. S8i**).

We next immunized mice with wild-type H2 HA or reoH2HA adjuvanted with alum and CpG oligodeoxynucleotide (**Fig. 4e**). We chose this combined alum/CpG adjuvant because we found in an immunization study with another HA (A/Shanghai/2/2013) that it further elevated the overall antibody titers compared to alum alone (**Fig. S9a,b**). After three injections, mice from both groups developed robust H2 HA-specific IgG responses (**Fig. 4f**). Despite similar antigen-specific titers, reoH2HA elicited much higher antibody titers against HA-stem, when we tested binding of antisera to the H1-stabilized stem (H1-SS) protein^29^ (**Fig. 4g**).

To test whether stem-binding translated to increased cross-reactivity, we measured binding of antisera to different group 1 and 2 HAs. After two doses, antisera from mice immunized with reoH2HA showed higher cross-reactive titers than those immunized with wild-type H2 HA (**Fig. S9c**). Even after three doses, mice immunized with H2 HA only cross-reacted with half of the HAs we tested (H1 CA/09, H5 VT/04, H7 SH/13), although with relatively low titers. In contrast, all mice immunized with reoH2HA developed cross-reactive responses against both group 1 and 2 HAs tested (**Fig. 4h**, **Fig. S9d**). To further evaluate if cross-reactive responses were focused, at least partially, on neutralizing epitopes of HA-stem, we analyzed whether antisera competed for binding with a known stem-specific bnAb (MEDI8852). While there was negligible competition with MEDI8852 in the H2 HA group (**Fig. 4i**), antisera against reoH2HA clearly competed with MEDI8852. As a control, antisera from neither group showed much competition for binding with another bnAb (FluA-20) that targets the trimer interface of HA (**Fig. S9e**). These results indicate that alum-bound reoH2HA directed antibody responses away from the head and toward the stem region of H2 HA.

## DISCUSSION

It remains challenging to acquire durable and protective immunity from vaccinations against viruses with high sequence variability, such as influenza and SARS-CoV-2. However, the discovery of bnAbs that target conserved epitopes on viral glycoproteins suggests the possibility of developing universal vaccines against these targets and has prompted the development of epitope-focused vaccine candidates to direct antibody responses toward bnAb epitopes^7^. In the case of influenza, intense efforts to focus antibody responses to HA-stem include cross-strain boosting^16–19^, mosaic display^20,21^, epitope resurfacing^22^, epitope masking^24–26^, and protein dissection^27–30^. While these diverse immunofocusing approaches have led to substantial progress toward broadening the breadth of the antibody response, it has not been possible to elicit an antibody response with a single immunogen that consistently cross-reacts with HAs from both group 1 and 2 influenza A subtypes.

Here, we introduce a new approach to immunofocusing – antigen reorientation. Our hypothesis was that the reorientation of antigens could alter epitope accessibility and redirect the immune response to less dominant but more desirable epitopes. Using influenza HA as a model, we reoriented the full-length HA ectodomain in an “upside-down” configuration facilitated by its binding to alum, to redirect antibody responses away from the variable HA-head and toward the highly conserved HA-stem. We envisioned that this approach could increase the exposure of the poorly accessible HA-stem that was originally occluded by HA-head on influenza virions^31,32^, while simultaneously sterically decreasing the exposure of HA-head – in the context of an intact HA trimer that preserves the native structure of HA-stem.

To test this, we genetically inserted oligoD into HA-head and found that this modification enhanced antigen-binding to alum in an “upside-down” configuration, likely through electrostatic interactions. Compared to existing immunofocusing approaches, our strategy simultaneously preserved the native structure and conformational epitopes of HA and mitigated the immunodominance of HA-head by tethering it to alum. Immunization with wild-type H2 HA on alum induced an antibody response that cross-reacted with several of the HAs tested, consistent with previous results^42^. By contrast, reoH2HA on alum elicited higher titers of cross-reactive antibodies against all group 1 and 2 HAs tested, in support of our hypothesis that antigen reorientation redirected antibody responses to the HA-stem region. Data from the competition assay with the HA-stem-directed bnAb MEDI8852 further indicate that antisera derived against reoH2HA include antibodies that target neutralizing epitopes on HA-stem, while antisera raised against wild-type H2 HA generally do not. Therefore, increased exposure and accessibility of HA-stem in the reoriented HA directed a fraction of the overall response to important stem epitopes.

Our results provide proof-of-concept that the reorientation of viral antigens leads to a stem-directed, cross-reactive antibody response. In future work, it will be important to explore higher doses or more potent adjuvants than those used here with the aim of further boosting cross-reactive antibody titers. As well, whether such cross-reactive antibody responses elicited by reoriented antigens protect against lethal challenges by direct viral neutralization or Fc-mediated mechanisms remains to be determined.

The simplicity of our approach makes it readily generalizable to other antigens since it only requires molecular cloning and screening of insertion sites. Meanwhile, existing methods for attaching antigens to alum require the synthesis of linker molecules, followed by chemical conjugation and subsequent purification steps^37,43^. Importantly, identifying sites where insertion of oligoD is permissive for protein expression in mammalian cells will likely be required to generate antigens with preserved native structures^44^. The trivalent anchoring afforded by three distinct oligoD motifs within each HA trimer rigidifies the antigen-alum binding interface and reduces the possibility of antigens “lying down” on alum. In addition, conceptually similar motifs that harness electrostatic interactions (*e.g.*, oligomeric glutamate residues) and other protein tags or binding partners^45–47^ may substitute or could be combined with oligoD for antigen reorientation and high-resolution immunofocusing purposes.

In summary, we introduce a generalizable immunofocusing strategy that enables the control of antigen orientation on alum by oligoD insertion. Our results showed consistent cross-reactivity toward group 1 and group 2 HAs with a single immunogen (reoH2HA) that encodes HA from a single influenza subtype. This approach has the potential to accelerate the development of epitope-focused vaccines against a broad range of viruses and other infectious agents.

## METHODS

### Cell lines

HEK-293T cells were purchased from American type culture collection (ATCC) and maintained in D10 medium – Dulbecco’s Modified Eagle Medium (DMEM, Corning) supplemented with 10% fetal bovine serum (GeminiBio) and 1% penicillin/streptomycin/*L*-glutamine (GeminiBio). HeLa-ACE2/TMPRSS2 cells were a generous gift from Dr. Jesse Bloom at the Fred Hutchinson Cancer Research Center and were maintained in D10 media.

Expi-293F cells were purchased from Thermo Fisher and maintained in Freestyle293/Expi-293 media (2:1, *v/v*, Thermo Fisher) in polycarbonate shaking flasks (Triforest Labware). Stellar^TM^ and BL21(DE3) competent cells were purchased from Takara Bio and Thermo Fisher, respectively.

### Antibodies

Monoclonal antibodies (mAbs) against Ebola glycoprotein (mAb114, c13C6, ADI-15742, KZ52 and ADI-16061), SARS-CoV-2 spike (COVA2-15, CB6 and CR3022), or influenza hemagglutinin (CH65, H2897, 6649, MEDI8852, CR9114, FI6v3, 8F8 and 8M2) were expressed in Expi-293F cells *via* transient transfection.

Mouse anti-His Tag antibody (clone J099B12, BioLegend #652502), goat anti-mouse IgG, HRP conjugated (clone Poly4053, BioLegend #405306), goat anti-mouse IgG1-HRP (SouthernBiotech #1070-05), goat anti-mouse IgG2a-HRP (SouthernBiotech #1083-05), goat anti-mouse IgG2b-HRP (SouthernBiotech #1093-05), or rabbit anti-human IgG, HRP conjugated (Abcam #ab6759) were used as detecting antibodies for Western-blot analysis or enzyme-linked immunosorbent assays (ELISAs).

For the analysis of germinal center responses, we stained cells with Ghost Dye Violet 510 (Tonbo Biosciences), anti-mouse CD16/32 (clone 2.4G2, BD Biosciences #553142), CD3 (clone 17A2, BioLegend #100216), CD4 (clone GK1.5, BioLegend #100469), CXCR5 (clone L138D7, BioLegend #145529), PD1 (clone 29F.1A12, BioLegend #135228), CD19 (clone 6D5, BioLegend #115534), CD95 (clone Jo2, BD Biosciences #557653) and CD38 (clone 90, BD Biosciences #740245).

### Antigen and antibody cloning

DNA encoding the Ebola glycoprotein (GP, Mayinga, Zaire, 1976) ectodomain with the mucin-like domain deleted (residues 1-308, 491-656) and the transmembrane domain replaced with a GCN4^48^ or foldon trimerization^49^ domain followed by an Avi-Tag^50^ and a hexahistidine tag was cloned into a mammalian expression vector (pADD2) by In-Fusion (Takara Bio). Similarly, DNAs encoding SARS-CoV-2 spike (residues 1–1143 from the Wuhan-Hu-1 genome sequence, GenBank MN9089473) or influenza hemagglutinins (HAs of H1N1 A/New Caledonia/20/99, H1N1 A/California/07/2009, H2N2 A/Japan/305/1957, H5N1 A/Vietnam/1203/2004, H3N2 A/Victoria/3/1975, H7N7 A/FPV/Dutch/1927, H7N9 A/Shanghai/2/2013) were cloned into the pADD2 vector with a GCN4 or foldon trimerization domain followed by an Avi-Tag and a hexahistidine tag on the C-terminus. DNA encoding Gen6’ H1-stabilized stem (H1-SS) was cloned into the pADD2 vector with a GCN4 trimerization domain followed by an Avi-Tag and a hexahistidine tag on the C-terminus. OligoD was inserted into the C-terminus after the hexahistidine tag in pADD2 expression plasmids. OligoD was also inserted into the flexible loop regions on Ebola GP or head regions of H2 HA (A/Japan/305/1957) in pADD2 expression plasmids. DNA encoding angiotensin-converting enzyme 2 – Fc fusion (ACE2-Fc) or ZsGreen with an Avi-tag, a hexahistidine tag and oligoD insertion to the C-terminus (ZsGreen-Avi-His-12D) was cloned into the pADD2 vector. Superfolder green fluorescent protein with a GCN4, an Avi-tag and a hexahistidine tag on the C-terminus (GFP-GCN4-Avi-His) was cloned into a pET28a bacterial expression vector^51^. DNA fragments encoding the variable heavy chain (HC) and light chain (LC) of antibodies were inserted into an expression plasmid encoding VRC01 HC and LC constant domains by In-Fusion.

All plasmid sequences were confirmed by Sanger sequencing (Sequetech). For transfection purposes, plasmids were transformed into Stellar^TM^ cells, isolated by Maxiprep kits (NucleoBond Xtra Maxi kit, Macherey Nagel), filtered through a sterile 0.45-µm membrane in a biosafety cabinet and stored at -20 °C.

### Protein expression and purification

All antigens and antibodies were expressed in Expi-293F cells. Expi-293F cells were cultured at 37 °C under constant shaking (120 rpm) in a humidified CO_2_ (8%) incubator. Expi-293F cells were transfected at a density of 3-4 × 10^6^ cells/mL. For 200 mL transfection of antigen proteins, the transfection mixture was made by adding 120 µg plasmid DNA (from Maxiprep) to 20 mL expression media, followed by the dropwise addition of 260 µL FectoPro transfection reagent (Polyplus) with vigorous mixing. For antibody production in 200 mL Expi-293F cells, the transfection mixture contained 60 µg LC plasmid DNA and 60 µg HC plasmid DNA. Transfection mixtures were incubated at room temperature for 10 minutes before being transferred to Expi-293F cells. *D*-glucose (4 g/L, Sigma-Aldrich) and valproic acid (3 mM, Acros Organics) were added to the cells immediately post-transfection to increase recombinant protein production. Cells were boosted again with *D*-glucose three days post-transfection and harvested on day four by centrifugation at 7000 ×*g* for five min. The supernatant was filtered through a 0.22-µm membrane for subsequent purification processes. Biotinylated proteins were expressed using the same protocol but with the presence of the BirA enzyme. For screening purposes, we expressed proteins in small scales (5–10 mL Expi-293F cells) and analyzed the supernatant with Western blots. We used mAb114 (0.5 µg/mL) or anti-His Tag antibody (1:4000) to detect Ebola GP or Influenza HAs, respectively.

Antigen proteins with hexahistidine tags were purified with HisPur^TM^ Ni-NTA resin (Thermo Fisher). Briefly, the filtered supernatant from Expi-293F cells was mixed with Ni-NTA resin (1 mL resin per liter supernatant) and incubated at 4 °C overnight. The mixture was then passed through a gravity-flow column, washed with 20 mM imidazole in HEPES buffer saline (HBS, 20 mM HEPES, pH 7.4, 150 mM NaCl), and then eluted with 250 mM imidazole in HBS. Elution was concentrated with centrifugal filters (30 or 50 kDa MWCO, Millipore Sigma) and buffer-exchanged to HBS for size-exclusion chromatography using a Superose^®^ 6 Increase 10/300 GL column (Cytiva) on an ÄKTA Protein Purification System (Cytiva). Peak fractions were pooled, concentrated, buffer-exchanged to HBS with 10% glycerol and filtered through a 0.22-µm membrane.

All antibodies and ACE2-Fc were purified with MabSelect PrismA protein A chromatography. Filtered supernatant from Expi-293F cells was directly applied to a MabSelect PrismA column (Cytiva) on an ÄKTA Protein Purification System. The column was washed with HBS and then antibodies were eluted with glycine (100 mM, pH 2.8) to HEPES buffer (1 M, pH 7.4). Fractions were concentrated and buffer-exchanged to HBS with 10% glycerol. Fragment antigen-binding (Fab) was prepared by digesting antibodies (IgG) with Lysyl endopeptidase (Wako Chemicals). After digestion, Fabs were purified with Protein-A Agarose (Thermo Fisher) and buffer-exchanged to HBS with 10% glycerol.

BL21(DE3) competent cells were transformed with the pET28a plasmid encoding GFP-GCN4-Avi-His and induced at an optical density of 0.6 with isopropyl *β*-*D*-*1*-thiogalactopyranoside (IPTG, 1 mM) for three hours at 37 °C. Cells were harvested by centrifugation at 8000 ×*g* for five min, resuspended in HBS with protease inhibitors (Halt™ Protease and Phosphatase Inhibitor Cocktail, Thermo Fisher) and lysed by sonication. Cell lysates were centrifuged at 16,000 ×*g* for 60 minutes and the supernatant was further purified with Ni-NTA resin, as described above.

The concentration of all proteins was determined by absorbance at 280 nm (A280), and the purity was assessed by protein gel electrophoresis. Protein samples were flash-frozen in liquid nitrogen and stored at -20 °C.

### Differential scanning fluorimetry

Thermal melting profiles of proteins were measured by differential scanning fluorimetry on a Prometheus NT.48 instrument (NanoTemper). Protein samples (0.1 mg/mL) were loaded into glass capillaries (NanoTemper) and then subjected to a temperature gradient from 20 to 95 °C at a heating rate of 1 °C per min. Alternatively, protein samples (0.1 mg/mL) were pre-mixed with alum (10 mg/mL, Alhydrogel^®^, InvivoGen) at a ratio of 1:10 (protein: alum, *w/w*) for 30 minutes at room temperature before loading into glass capillaries. HBS and alum (diluted to 1 mg/mL in HBS) were also loaded into glass capillaries and measured as controls. Intrinsic fluorescence (350 nm and 330 nm) was recorded as a function of temperature. Thermal melting curves were plotted using the first derivative of the ratio (350 nm/330 nm). Melting temperatures were calculated automatically by the instrument (PR.ThermControl software, version 2.3.1) and represented peaks in the thermal melting curves.

### Biolayer interferometry (BLI)

BLI experiments were performed on an OctetRed 96 system (FortéBio). All samples were diluted with the Octet buffer (DPBS with 0.05% Tween-20 and 0.1% BSA), and assays were performed under agitation (1000 rpm). mAbs (200 nM) were loaded onto anti-human IgG Fc capture (AHC) Biosensors (FortéBio) and then dipped into antigen solutions (150–200 nM) for binding analysis, followed by dissociation into the Octet buffer. Alternatively, biotinylated antigens (200 nM) were loaded onto Streptavidin biosensors (Sartorius) and then dipped into antibody solutions (200 nM) for binding analysis, followed by dissociation into the Octet buffer. Serial dilutions of Fabs were used to measure the affinity (K_D_) for biotinylated antigens. Data were processed by Data Analysis software (version 9.0.0.15, FortéBio) and then plotted. Shifts in nanometers of mAb-binding to wild-type proteins were used as the maximal binding value (1.0) and shifts from oligoD-modified proteins were normalized as a fraction of the maximal binding value.

### Antigen-alum binding assays

Wild-type GP or oligoD-modified GP were first incubated with alum (protein: alum, 1:10, *w/w*) for 30 minutes in PBS at room temperature and then naïve mouse serum was added to the mixture to a final concentration of 10% (*v/v*). The mixture with alum alone or wild-type GP alone served as controls. Mixtures were further incubated at 37 °C under constant shaking (220 rpm) on an orbital shaker for 24 hours before being centrifuged to pellet alum (10,000 ×*g* for five min). The supernatant was collected for ELISA to measure the concentration of unbound protein antigens. Alum pellets were rinsed extensively with PBS and re-pelleted again (twice) by centrifugation, then resuspended in SDS-PAGE sample loading buffer for Western-blot analysis to determine the amount of alum-bound protein.

To measure the amount of unbound GP, we coated Nunc 96-well MaxiSorp plates (Thermo Fisher) with mAb114 (2 µg/mL in DPBS, 50 µL per well) for one hour at room temperature. Plates were washed three times with Milli-Q H_2_O (300 µL) using a plate washer (ELx405 BioTek) and then blocked with ChonBlock (100 µL per well, Chondrex) overnight at 4 °C. For subsequent steps, all dilutions were made in DPBS with 0.05% Tween-20 and 0.1% BSA, and ELISA plates were rinsed with PBST (PBS with 0.1% Tween 20, 300 µL, three times) in between steps. Antigens with proper dilutions were added to the plate and detected by the mouse anti-His tag antibody (1:4000). Goat anti-mouse IgG, HRP-conjugated (1:4000) was then added for one-hour incubation before rinsing with PBST six times. ELISA plates were developed with the TMB (3,3’,5,5’-tetramethylbenzidine) substrate (1-Step Turbo-TMB, Thermo Fisher) for six minutes and terminated with sulfuric acid (2M). Absorbance at 450 nm was recorded on a microplate reader (Synergy^TM^ HT, BioTek). GP with a series of known concentrations was used to establish a standard curve, and the amount of antigen was quantified by fitting the absorbance values to this curve.

For Western-blot analysis, alum pellets were resuspended in sample loading buffer and boiled at 95 °C for ten minutes before being applied to a precast polyacrylamide gel (4–20% Mini-PROTEAN^®^, Bio-Rad). Proteins were then transferred to a nitrocellulose membrane using the Trans-Blot Turbo transfer system (Bio-Rad). Blots were blocked in PBST with 10% non-fat dry milk (Bio-Rad), incubated with mAb114 (0.5 µg/mL in PBST with 10% non-fat dry milk) and then detected with rabbit anti-human IgG, HRP-conjugated (1:4000 in PBST with 10% non-fat dry milk). Luminescent signals were developed using a luminol-based substrate (Pierce ECL, Thermo Scientific) for imaging on a chemiluminescence imager (GE Amersham Imager 600).

### Streptavidin-and alum-based ELISA

To investigate antigen orientation on alum, we measured mAb binding in streptavidin-or alum-based ELISAs. For streptavidin-based ELISAs, Nunc 96-well MaxiSorp plates were coated with streptavidin (4 µg/mL in DPBS, 60 µL per well) for one hour at room temperature. These plates were washed three times with Milli-Q H_2_O and then blocked with ChonBlock (120 µL per well) overnight at 4 °C. For subsequent steps, all dilutions were made in DPBS with 0.05% Tween-20 and 0.1% BSA, and ELISA plates were rinsed with PBST in between steps. Biotinylated H2 HA-12D or reoH2HA (2 µg/mL) were added to the plates and incubated for one hour at room temperature. mAbs (8F8, 8M2, MEDI8852 and FI6v3) were serially diluted (10-fold dilution starting from 20 nM) and then added to the ELISA plates for one-hour incubation at room temperature. Rabbit anti-human IgG, HRP-conjugated (1:4,000) was added for one-hour incubation before rinsing with PBST six times.

For alum-based ELISA, Nunc 96-well MaxiSorp plates were coated with ZsGreen-Avi-His-12D (abbreviated as ZsGreen-12D, 4 µg/mL in DPBS, 60 µL per well) for one hour at room temperature. These plates were washed three times with Milli-Q H_2_O and then blocked with ChonBlock overnight at 4 °C. For subsequent steps, all dilutions were made in DPBS with 0.05% Tween-20 and 0.1% BSA unless otherwise noted, and ELISA plates were rinsed with PBST in between steps. Alum (100 µg/mL in HBS) was added to the plates and incubated for one hour at room temperature. After rinsing, H2 HA-12D or reoH2HA (2 µg/mL) was added to the plates and incubated for one hour at room temperature. mAbs were serially diluted (10-fold dilution starting from 20 nM) and then added to the ELISA plates for one-hour incubation at room temperature. Rabbit anti-human IgG, HRP-conjugated (1:4,000) was added for one-hour incubation before rinsing with PBST six times. ELISA plates were developed with the TMB substrate for five minutes and terminated with sulfuric acid. Absorbance at 450 nm was recorded on a microplate reader.

### Mouse immunization studies

All animals were maintained in accordance with the Public Health Service Policy for “Human Care and Use of Laboratory Animals” under a protocol approved by the Stanford University Administrative Panel on Laboratory Animal Care (APLAC-33709). Female BALB/c mice (6-8 weeks) were purchased from Jackson Laboratories and randomly grouped for the studies. Antigens were mixed with adjuvants before injections, with the total volume adjusted to 100 µL with HBS. Injection mixtures were incubated at room temperature for 30 minutes before use.

To compare GP-12D with wild-type GP, we performed a single-dose immunization in two groups of BALB/c mice (*n*=6) with 5 µg protein antigens adjuvanted with 150 µg alum *via* subcutaneous injection. To compare spike-12D with wild-type spike, we immunized two groups of BALB/c mice (*n*=10) with 5 µg protein antigens adjuvanted with 150 µg alum *via* subcutaneous injection on days 0, 21, and 49. To compare H1 HA-12D with wild-type H1 HA, we immunized two groups of BALB/c mice (*n*=10) with 5 µg protein antigens adjuvanted with 150 µg alum *via* subcutaneous injection on days 0 and 56. To examine the effect of CpG oligodeoxynucleotide, we immunized two groups of BALB/c mice (*n*=5) with 5 µg H7 HA-12D adjuvanted with 150 µg alum alone or 150 µg alum and 1 µg CpG (ODN 1826, InvivoGen) *via* subcutaneous injection on days 0 and 21. And to compare reoH2HA with wild-type H2 HA, we immunized two groups of BALB/c mice (*n*=5) with 5 µg protein antigens adjuvanted with 150 µg alum and 1 µg CpG *via* subcutaneous injection on days 0, 21 and 70.

Pre-immune, interim and final blood samples were collected by retro-orbital bleeding into serum gel tubes (Sarstedt). Serum gel tubes were centrifuged at 10,000 ×*g* for 6 min, and sera were collected and stored at -80 °C.

### Serum ELISAs

We used proteins with a different trimerization domain from the immunogens for ELISA binding analysis of antisera. Nunc 96-well MaxiSorp plates were coated with streptavidin for one hour at room temperature. These plates were washed three times with Milli-Q H_2_O using a plate washer and then blocked with ChonBlock overnight at 4 °C. For subsequent steps, all dilutions were made in DPBS with 0.05% Tween-20 and 0.1% BSA, and ELISA plates were rinsed with PBST in between steps. Biotinylated antigens (2 µg/mL) were added to the plates and incubated for one hour at room temperature. Mouse antisera were serially diluted (5-fold dilution) and then added to the ELISA plates for one-hour incubation at room temperature.

Goat anti-mouse IgG, HRP-conjugated (1:4000) was added for one-hour incubation before rinsing with PBST six times. ELISA plates were developed with the TMB substrate for six minutes and terminated with sulfuric acid. Absorbance at 450 nm was recorded on a microplate reader.

To test the cross-reactivity toward different HAs, streptavidin-coated ELISA plates were incubated with biotinylated HAs (2 µg/mL) from the following strains: A/New Caledonia/20/1999 (H1 NC/99), A/California/07/2009 (H1 CA/09), A/Japan/305/1957 (H2 JP/57), A/Vietnam/1203/2004 (H5 VT/04), A/Victoria/3/1975 (H3 VC/75), A/FPV/Dutch/1927 (H7 NT/27), A/Shanghai/2/2013 (H7 SH/13).

For competition ELISA assays, competing antibodies (*e.g.*, MEDI8852 or FluA-20, 100 nM) were preincubated on the plate with antigens for one hour before the addition of serially diluted mouse antisera.

### Pseudotyped lentiviruses and neutralization assays

Ebola GP-pseudotyped lentiviruses (EBOV) encoding a luciferase-ZsGreen reporter were produced in HEK-293T cells by co-transfection of five plasmids^52^. This five-plasmid system includes a packaging vector (pHAGE-Luc2-IRES-ZsGreen), a plasmid encoding full-length Ebola GP (pCDNA3.1 EBOV FL-GP) and three helper plasmids (pHDM-Hgpm2, pHDM-Tat1b and pRC-CMV_Rev1b). One day before transfection, HEK-293T cells were seeded in 10-cm TC-treated culture dishes (5 × 10^6^ cells per culture dish, Corning). Transfection mixture was prepared by adding five plasmids (10 µg packaging vector, 3.4 µg GP-encoding plasmid and 2.2 µg of each helper plasmid) to 1 mL D10 medium, followed by the dropwise addition of BioT transfection reagent (30 µL, Bioland Scientific) with vigorous mixing. After 10-minute incubation at room temperature, the transfection mixture was transferred to HEK-293T cells in the culture dish. Culture medium was replenished 24 hours post-transfection, and after another 48 hr, viruses were harvested and filtered through a 0.45-µm membrane. Pseudotyped lentiviruses were aliquoted, flash-frozen in liquid nitrogen, stored at -80 °C, and titrated before further use.

SARS-CoV-2 spike-pseudotyped lentiviruses encoding a luciferase-ZsGreen reporter were produced using the same method but with plasmids encoding SARS-CoV-2 spike (HDM-SARS2-spike-delta21, Addgene #155130) or its variants of concern. Wild-type (WT) pseudovirus included a D614G mutation. B.1.351 (Beta) pseudovirus included D80A, D215G, Δ240-242, K417N, E484K, N501Y, D614G and A701V mutations. B.1.617.2 (Delta) pseudovirus included T19R, T95I, G142D, Δ156-157, R158G, L452R, T478K, D614G, P681R and D950N mutations. B.1.617.2/AY1 (Delta plus) pseudovirus included V70F, A222V, W258L and K417N mutations in addition to the ones in B.1.617.2. B.1.529.1 (Omicron BA.1) pseudovirus included A67V, Δ69-70, T95I, G142D, Δ143-145, Δ211, L212I, G339D, S371L, S373P, S375F, K417N, N440K, G446S, S477N, T478K, E484A, Q493R, G496S, Q498R, N501Y, Y505H, T547K, D614G, H655Y, N679K, P681H, N764K, D796Y, N856K, Q954H, N969K and L981F mutations. B.1.529.2 (Omicron BA.2) pseudovirus included T19I, Δ24-26, A27S, G142D, V213G, G339D, S371F, S373P, S375F, T376A, D405N, R408S, K417N, N440K, S477N, T478K, E484A, Q493R, Q498R, N501Y, Y505H, D614G, H655Y, N679K, P681H, N764K, D796Y, Q954H and N969K mutations.

### Neutralization assays with mAbs

Neutralization of EBOV was validated with five Ebola GP-specific mAbs. HEK-293T cells were seeded in white-walled, clear-bottom 96-well plates (Thermo Fisher or Greiner Bio-One) at a density of 20,000 cells per well one day before the assay (day 0). On day 1, GP-specific mAbs (2 µM in HBS with 10% glycerol) were filtered with 0.22-μm sterile membranes and diluted with D10 media to 20 nM as the starting concentration. Subsequently, mAbs were serially diluted (10-fold dilution) in D10 media and mixed with EBOV (diluted in D10 medium, supplemented with polybrene, 1:1000, *v/v*) for one hour before being transferred to HEK-293T cells. On day 4, medium was removed, and 100 µL of luciferase substrates (BriteLite Plus, Perkin Elmer) were added to each well. Luminescent signals were recorded on a microplate reader (BioTek Synergy™ HT or Tecan M200). Percent infection was normalized to cells only (0% infection) and virus only (100% infection) on each plate. Neutralization assays were performed in biological duplicates with technical duplicates. Similarly, neutralization of SARS-CoV-2 pseudoviruses was validated with ACE2-Fc or three spike-specific mAbs. HeLa-ACE2/TMPRSS2 cells were seeded at a density of 6,000 cells per well one day before the assay (day 0). On day 1, ACE2-Fc or spike-specific mAbs (2 µM in HBS with 10% glycerol) were filtered and diluted to 20 nM as the starting concentration. Subsequently, ACE2-Fc or mAbs were serially diluted (10-fold dilution) in D10 media and mixed with SARS-CoV-2 pseudoviruses for one hour before being transferred to HeLa-ACE2/TMPRSS2 cells. On day 3, medium was removed, and luciferase substrates were added to each well. Luminescent signals were recorded on a microplate reader and converted to percent infection for subsequent analysis. Neutralization assays were performed in biological duplicates with technical duplicates.

### Serum neutralization assays

Antisera were heat-inactivated (56 °C, 30-60 min) before neutralization assays. Neutralization against EBOV was analyzed in HEK-293T cells with mouse antisera against Ebola GP. Neutralization against SARS-CoV-2 pseudoviruses was analyzed in HeLa-ACE2/TMPRSS2 cells with mouse antisera against spike. Briefly, cells were seeded in white-walled, clear-bottom 96-well plates one day before the assay (day 0). On day 1, antisera were serially diluted in D10 media and mixed with pseudoviruses for one hour before being transferred to cells. Assays against EBOV or SARS-CoV-2 pseudoviruses were read out with luciferase substrates three or two days post-infection, respectively. Percent infection was normalized on each plate. Neutralization titers (NT_50_) were calculated as the serum dilution, where a 50% inhibition of infection was observed. Neutralization assays were performed in technical duplicates.

### Flow cytometry analysis of germinal center (GC) responses

To study the germinal center responses, we immunized BALB/c mice (*n*=10 mice per group) with 5 µg protein antigens (GP or GP-12D) adjuvanted with 150 µg alum *via* subcutaneous injection on days 0, 7, and 14 (two groups per time point). On day 21, all six groups of mice and another group of naïve mice were euthanized for tissue collection. Draining lymph nodes were collected and triturated through a 70-μm cell strainer to make single-cell suspensions. Cells were then stained for viability with Ghost Dye Violet 510 for 5 minutes on ice in PBS with EDTA (2 mM). After rinsing, cells were blocked with Fc receptor antibody (anti-CD16/32) for 5 minutes on ice before staining with fluorescently conjugated antibodies in FACS staining buffer (PBS, 3% fetal bovine serum, 1 mM EDTA): CD3 (1:50), CD4 (1:200), CXCR5 (1:50), PD1 (1:200), CD19 (1:200), CD95 (1:200), CD38 (1:200). Surface staining was carried out for 30 minutes on ice and 15 minutes at 37 °C, followed by fixation for 10 minutes at room temperature. Data were acquired on a BD LSR-II flow cytometer and analyzed with FlowJo (version 10.4.0).

### Statistical analyses

Statistics were analyzed using GraphPad Prism software (version 9.3.1). Non-transformed data are presented as mean ± standard deviation (s.d.). Log-transformed data (ELISA titers and NT_50_) are presented as geometric mean ± s.d. Comparisons of two groups were performed using the two-tailed Mann–Whitney U test. Comparisons of more than two groups were performed using one-way ANOVA followed by a Bonferroni test. Comparisons of IgG titers or NT_50_ over time were performed using two-way ANOVA followed by a Bonferroni test. *P* values of 0.05 or less are considered significant and plotted.

### Data Availability

All data supporting the results in this study are available within the main text and its supplementary information. Additional data supporting the results of the study are also available from the corresponding authors on reasonable request.

## Acknowledgments

We thank members of the Kim Lab for fruitful discussions on the project. We thank Dr. Brian L. Hie and Dr. Mirella Bucci for comments on the manuscript. D.X. acknowledges the postdoctoral fellowship from the Stanford Maternal and Child Health Research Institute. S.T. acknowledges NIH NICHD grant K99HD104924 and the Merck fellowship from the Damon Runyon Cancer Research Foundation. This work was supported by an NIH Director’s Pioneer Award (DP1-AI158125), the Virginia & D.K. Ludwig Fund for Cancer Research, and the Chan Zuckerberg Biohub to P.S.K.

## Author contributions

D.X. and P.S.K. conceptualized the project, designed experiments, and analyzed data. D.X. and

C.L. performed the analysis of germinal center responses with research supervision from B.P.

D.X. and A.U. designed, expressed, and characterized protein antigens. P.A.B.W. provided suggestions on potential oligoD insertion sites on HA-head. P.A.B.W. and S.T. provided key reagents for influenza studies. D.X. and M.S. performed animal immunization studies and analysis of the antisera. D.X. and P.S.K. wrote the manuscript with input from all authors.

## SUPPLEMENTARY FIGURE AND LEGENDS

**Figure S1.**
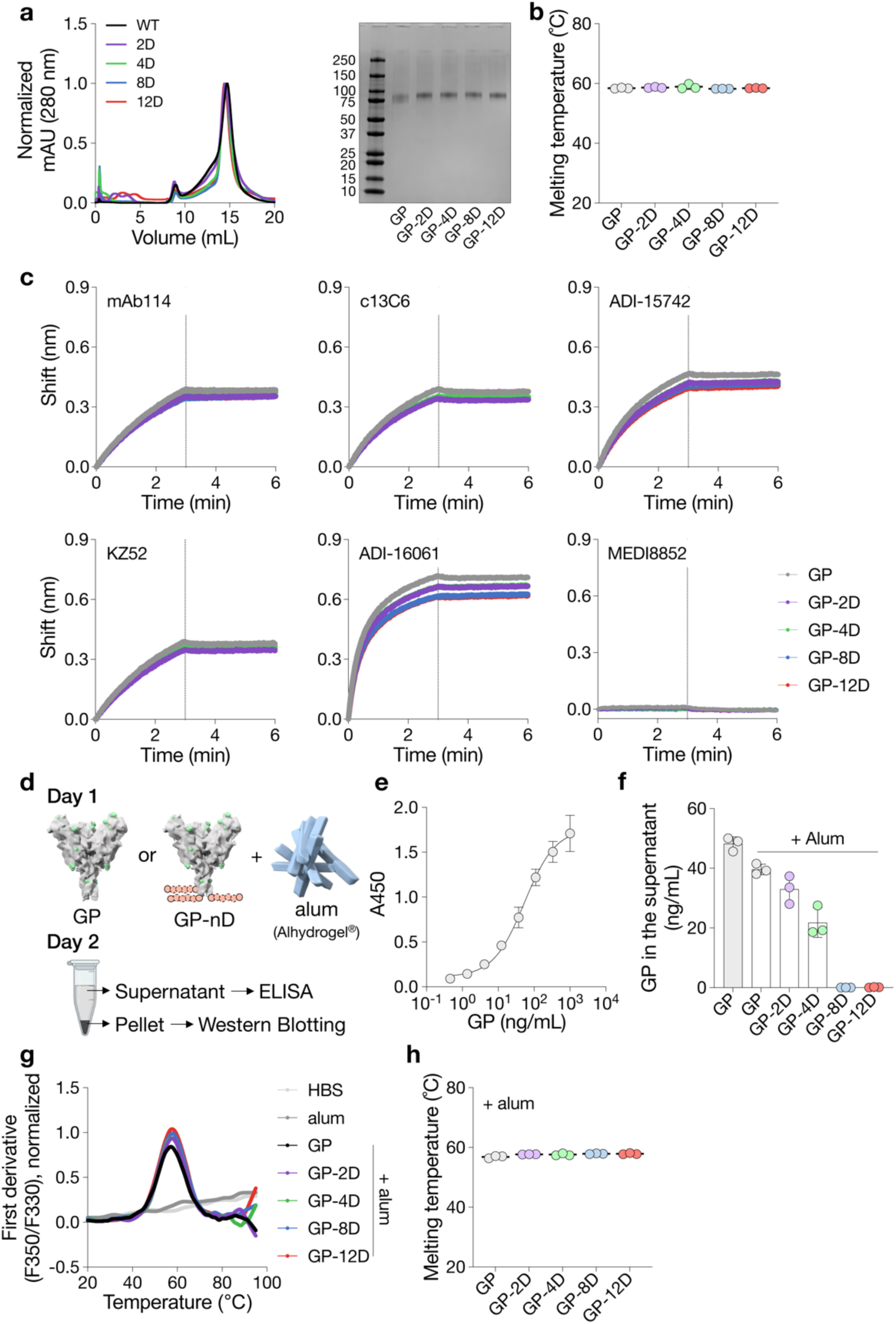
OligoD insertion into Ebola GP. **a**, Size-exclusion purification and gel electrophoresis of wild-type and oligoD-modified GP. Gel was stained with Coomassie brilliant blue. **b**, Thermal melting temperature (T_m_) of wild-type and oligoD-modified GP. Center values represent the mean (*n*=3 samples per group). **c**, BLI binding profiles of wild-type or oligoD-modified GP with five GP-specific mAbs (mAb114, c13C6, ADI-15742, KZ52 and ADI-16061). An HA-specific mAb (MEDI8852) served as a negative control. Vertical dashed lines indicate the beginning of dissociation. **d**, Alum-binding assay for wild-type or oligoD-modified GP. GP proteins were pre-mixed with alum for one hr at room temperature and then incubated in PBS containing naïve mouse serum for 24 hr at 37 °C. Upon centrifugation, the concentrations of unbound GP in the supernatant were quantified by ELISA. The rinsed pellet was analyzed by Western blotting to detect alum-bound GP. **e**, Standard curve established with known concentrations of GP for reference. Data are presented as mean ± s.d. (*n*=3 samples per group). **f**, Concentrations of unbound GP in the supernatant of GP-alum mixtures. Data are presented as mean ± s.d. (*n*=3 samples per group). **g,h**, Thermal melting profiles (**g**) and T_m_ (**h**) of wild-type or oligoD-modified GP in the presence of alum. Center values represent the mean (*n*=3 samples per group).

**Figure S2.**
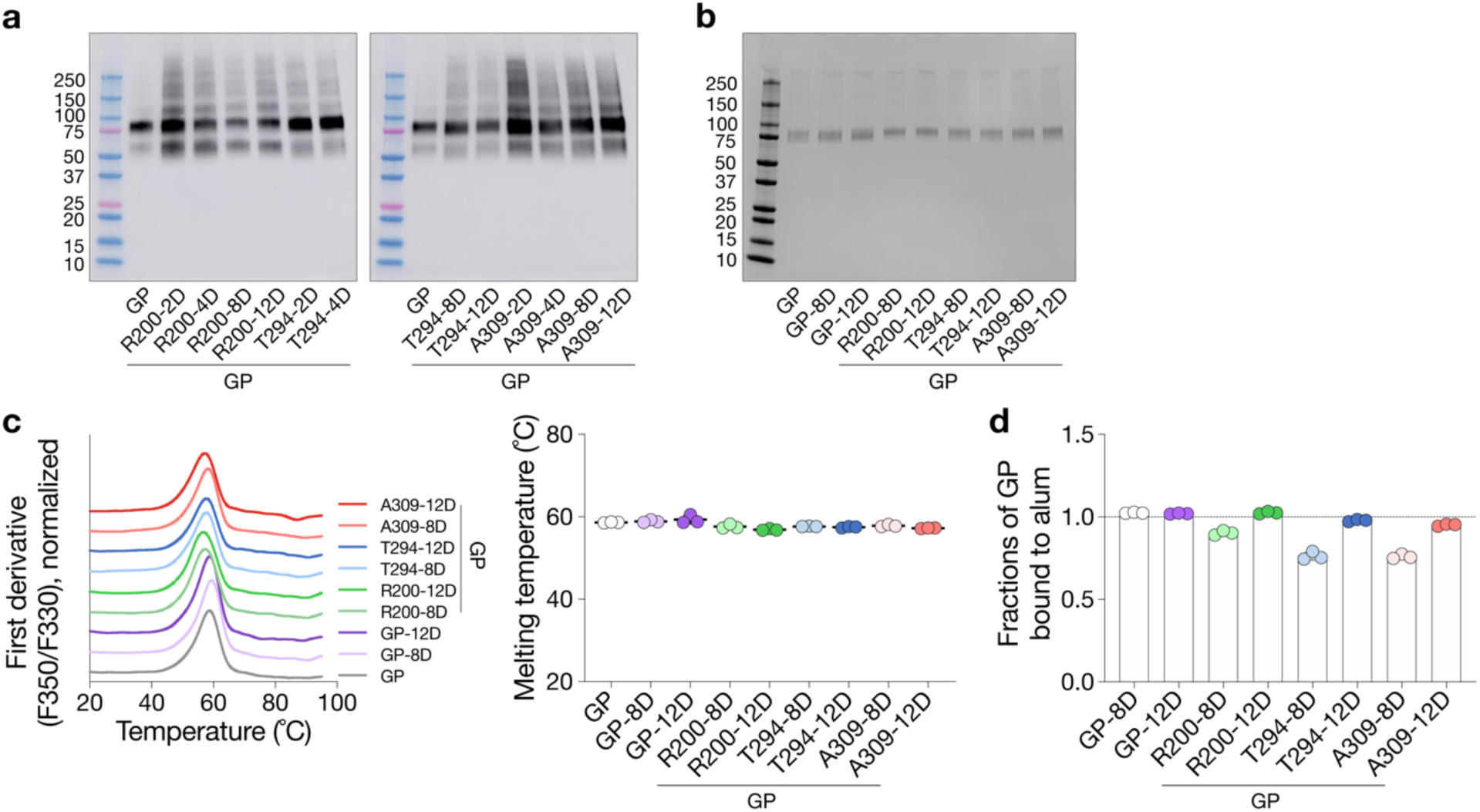
OligoD insertion into flexible loop regions on GP. **a**, Screening of oligoD insertion into flexible loop regions on Ebola GP by Western-blot analysis. Insertions (2,4,8 or 12D) were made after residues R200, T294 or A309 as indicated. Blots were detected by mAb114. **b**, Gel electrophoresis of oligoD-modified GP after purification. Gel was stained with Coomassie brilliant blue. **c**, Thermal melting profiles and T_m_ of wild-type and oligoD-modified GP. Center values in the dot plot represent the mean (*n*=3 samples per group). **d**, Quantification of the alum-bound fraction of oligoD-modified GP by ELISA (*n*=3 samples per group). Dashed line indicates 100% binding to alum.

**Figure S3.**
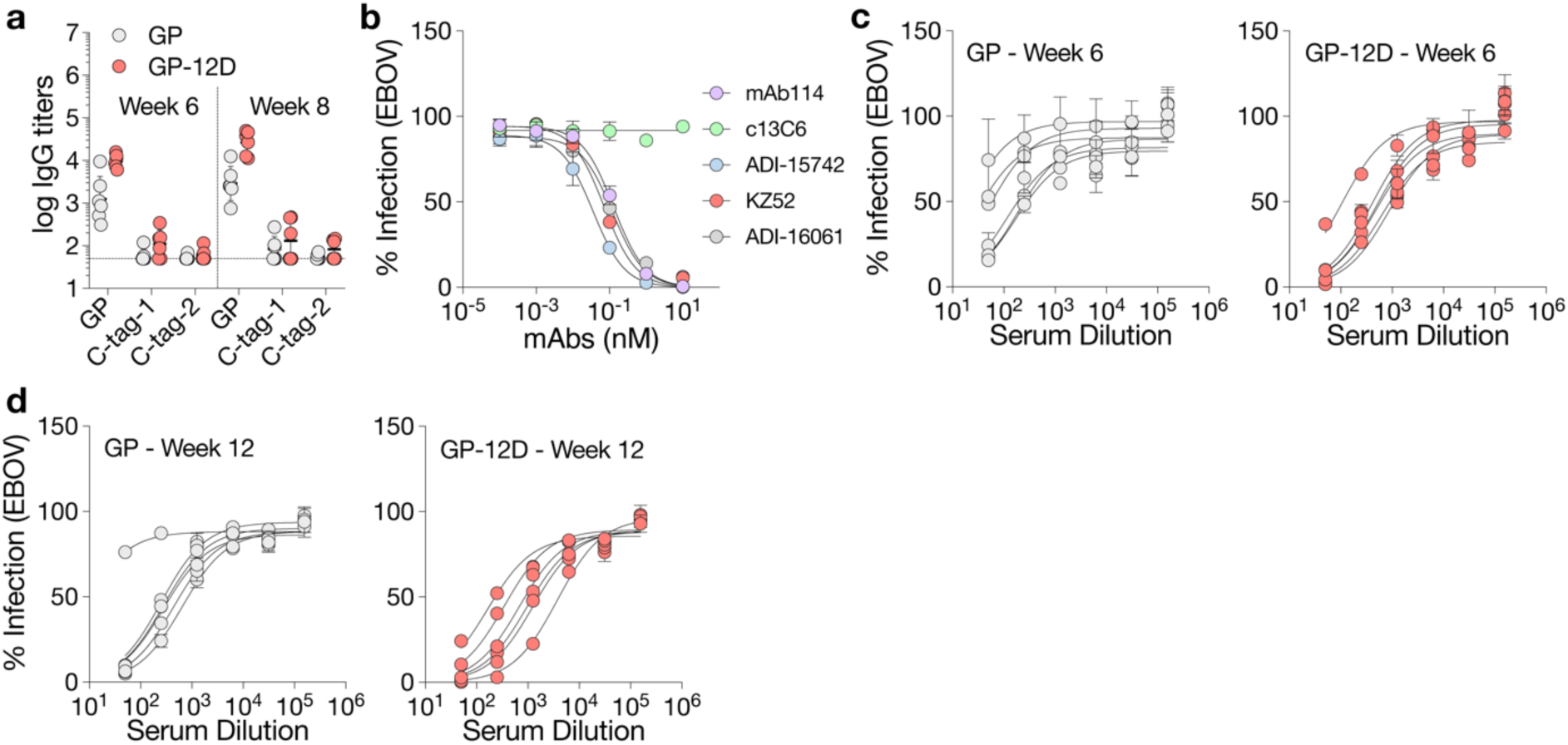
Immunogenicity of oligoD-modified GP. **a**, Serum binding titers of GP or C-terminal tags (C-Tag-1 = ZsGreen-Avi-His-12D; C-Tag-2 = GFP-GCN4-His) by ELISA. Dashed line indicates the limit of quantification. Data are presented as the geometric mean ± s.d. of the log-transformed values (*n*=6 mice per group). **b**, Validation of the neutralization assay against Ebola GP-pseudotyped lentiviruses (EBOV) with the five GP-specific mAbs. c13C6 is known to be non-neutralizing and serves as a control. Data are presented as mean ± s.d. (*n*=2 biological replicates with technical duplicates). **c,d**, Neutralization of Ebola GP-pseudotyped lentiviruses by antisera collected at six (**c**) or 12 weeks (**d**) post-immunization (*n*=6 mice per group). Each curve derived from a single mouse. Data are presented as mean ± s.d. of technical duplicates.

**Figure S4.**
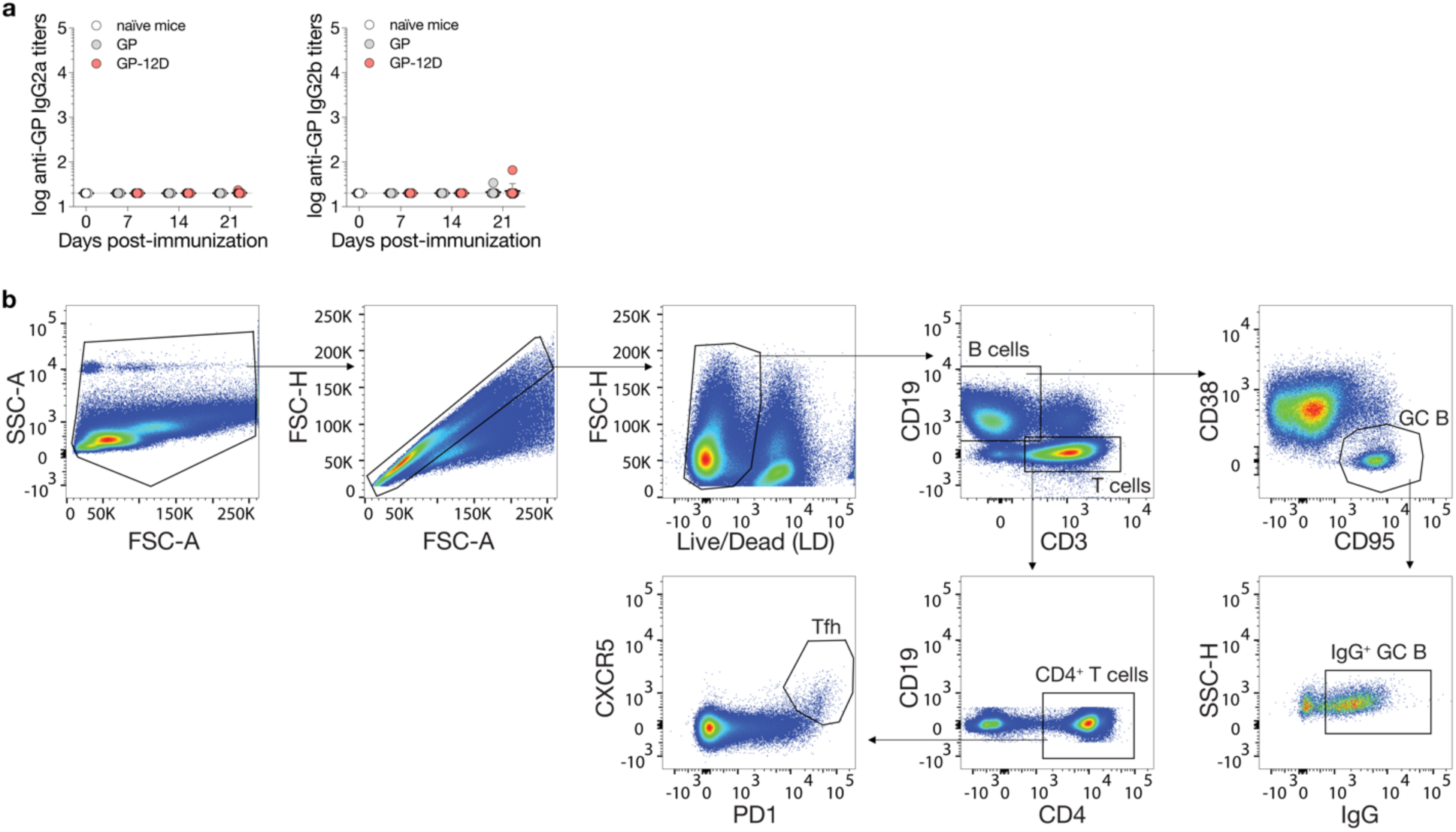
Analysis of germinal center responses. **a**, Serum GP-specific IgG2a or IgG2b titers 7, 14 or 21 days post-immunization (*n*=10 mice per antigen per time point). Dashed lines indicate the limit of quantification. Data are presented as the geometric mean ± s.d. of the log-transformed values. **b**, Gating strategy for the analysis of germinal center B cells (CD19^+^CD95^+^CD38^−^), IgG^+^ germinal center B cells and T follicular helper cells (CD3^+^CD4^+^PD1^+^CXCR5^+^) in the draining lymph nodes.

**Figure S5.**
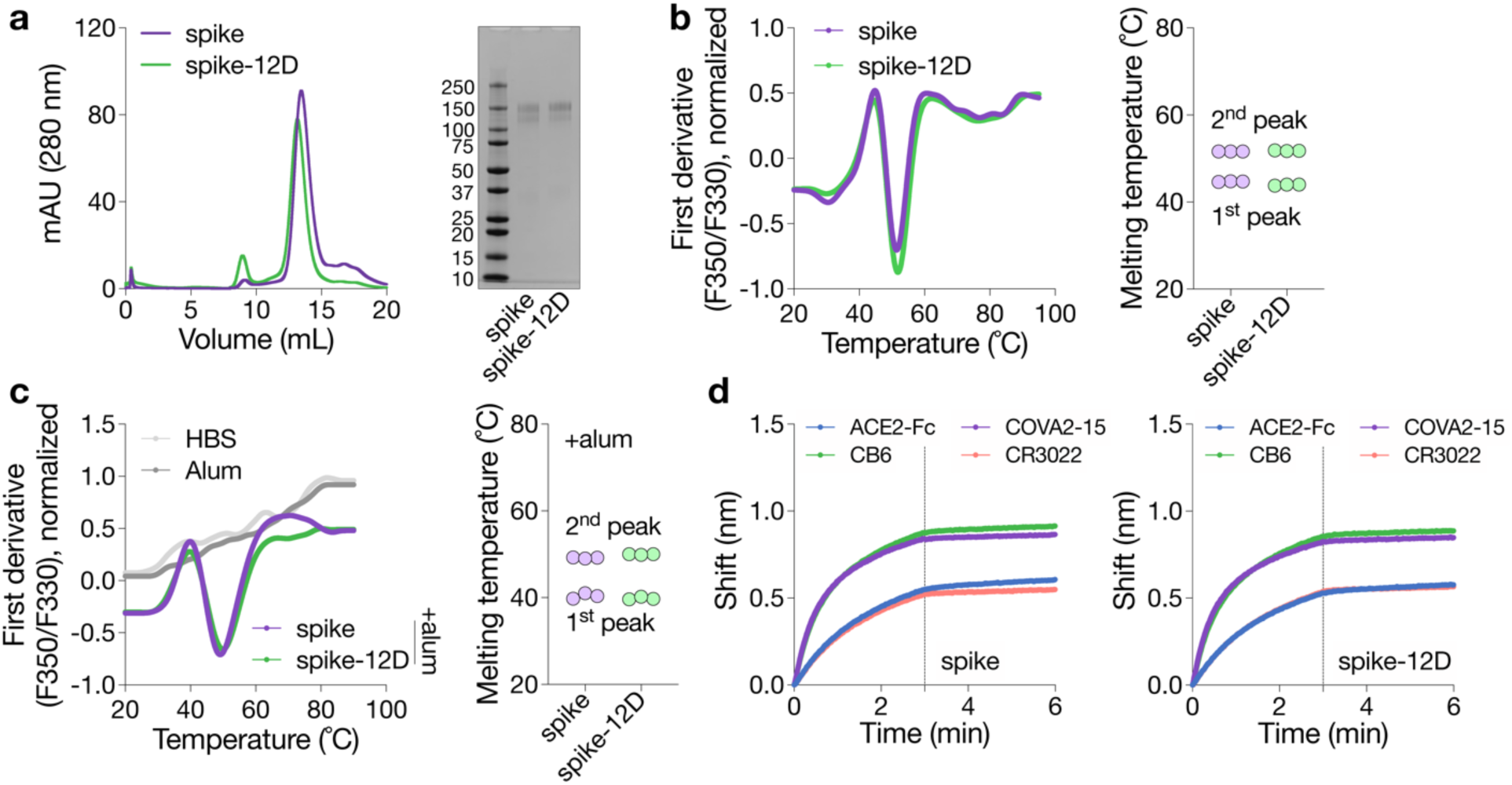
OligoD insertion into SARS-CoV-2 spike. **a**, Size-exclusion purification and gel electrophoresis of wild-type or oligoD-modified Spike. Gel was stained with Coomassie brilliant blue. **b**, Thermal melting profiles and T_m_ of wild-type and oligoD-modified spike (*n*=3 samples per group). **c**, Thermal melting profiles and T_m_ of wild-type and oligoD-modified spike in the presence of alum (spike: alum = 1:10, *w/w*) (*n*=3 samples per group). **d**, BLI binding profiles of wild-type or oligoD-modified spike with ACE2-Fc and three Spike-specific mAbs (COVA2-15, CB6 and CR3022). Dashed lines indicate the beginning of dissociation.

**Figure S6.**
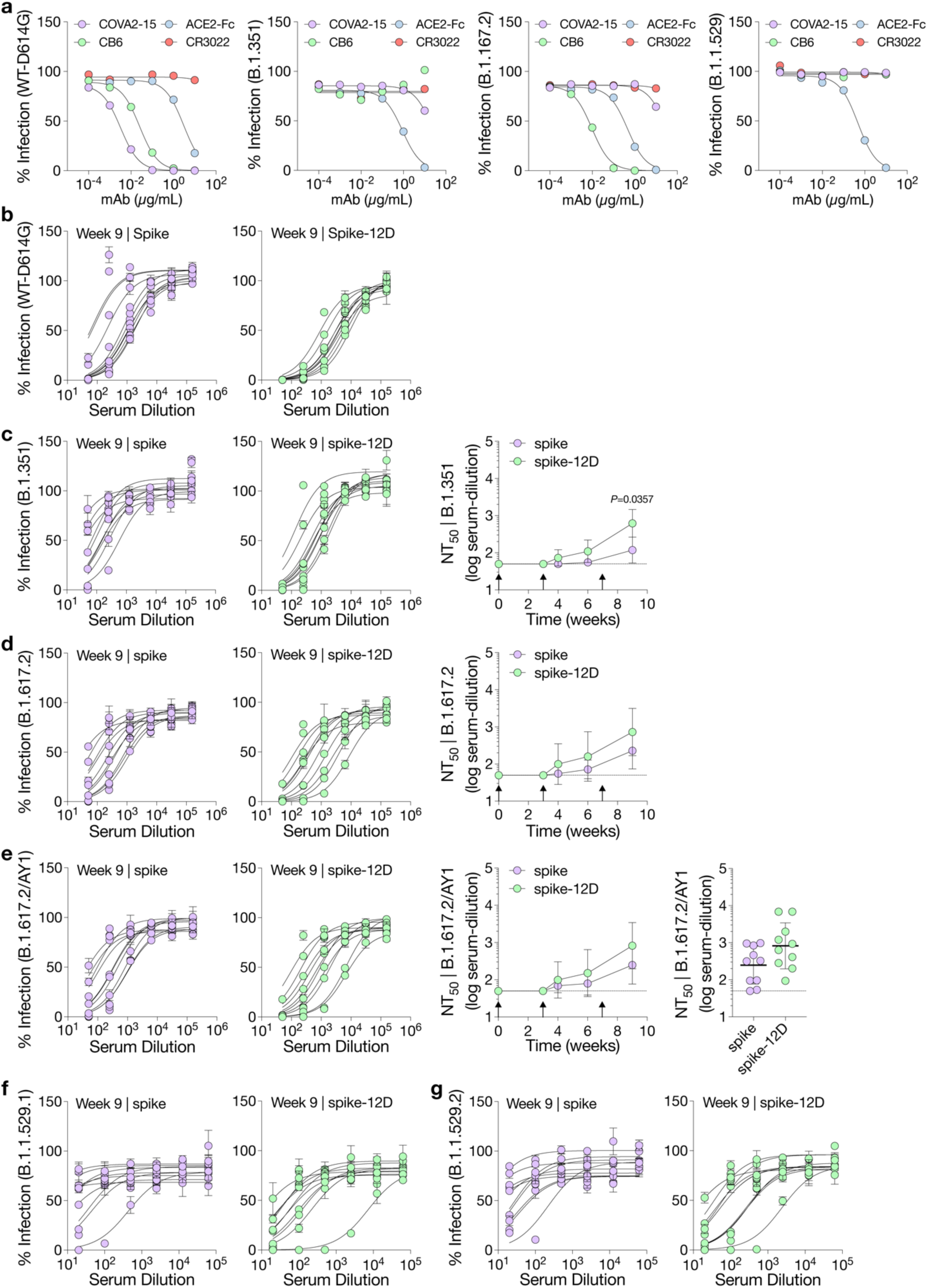
Neutralization of SARS-CoV-2 pseudoviruses. **a**, Validation of the neutralization assay with ACE2-Fc and three Spike-specific mAbs (COVA-215, CB6 and CR3022) against spike-pseudotyped lentiviruses and three variants of concerns (B.1.351, B.1.167.2 and B.1.1.529). Data are presented as mean ± s.d. (*n*=2 biological replicates with technical duplicates). **b-g**, Serum neutralization of wild-type SARS-CoV-2 pseudoviruses (**b**) and variants of concern, including B.1.351 (**c**), B.1.617.2 (**d**), B.1.617.2/AY1 (**e**), B.1.1.529.1 (**f**), and B.1.1.529.2 (**g**) (*n*=10 mice per group). Dashed lines indicate the limit of quantification of NT_50_ in **c-e**. Data are presented as mean ± s.d. of technical duplicates in neutralization curves. Data are presented as the geometric mean ± s.d. of the log-transformed values in NT_50_. Comparison of NT_50_ over time was performed using two-way ANOVA followed by a Bonferroni test. Comparison of two groups was performed using the two-tailed Mann–Whitney U test. *P* values of 0.05 or less were considered significant.

**Figure S7.**
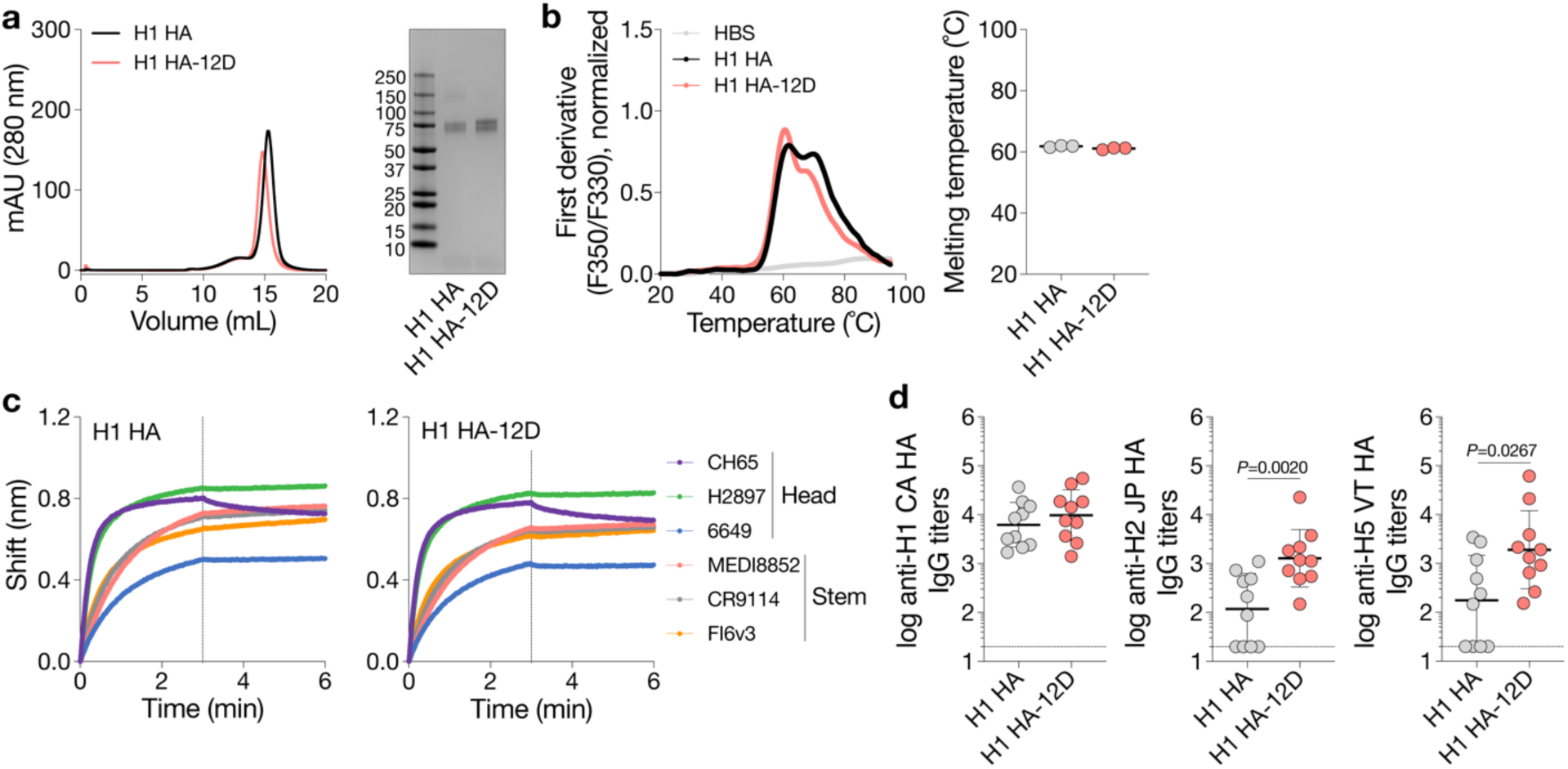
OligoD insertion into H1 HA. **a**, Size-exclusion purification and gel electrophoresis of wild-type or oligoD-modified H1 HA. Gel was stained with Coomassie brilliant blue. **b**, Thermal melting profiles and T_m_ of wild-type and oligoD-modified H1 HA. Center values represent the mean (*n*=3 samples per group). **c**, BLI binding profiles of wild-type or oligoD-modified H1 HA with three HA-head-directed mAbs (CH65, H2897 and 6649) and three HA-stem-directed mAbs (MEDI8852, CR9114 and FI6v3). Dashed lines indicate the beginning of dissociation. **d**, Serum binding titers to different group 1 HAs (H1 CA – A/California/07/2009, H2 JP – A/Japan/305/1957 and H5 VT – A/Vietnam/1203/2004). (*n*=10 mice per group). Dashed lines indicate the limit of quantification. Data are presented as the geometric mean ± s.d. of the log-transformed values. Comparison of two groups was performed using the two-tailed Mann– Whitney U test. *P* values of 0.05 or less were considered significant and plotted.

**Figure S8.**
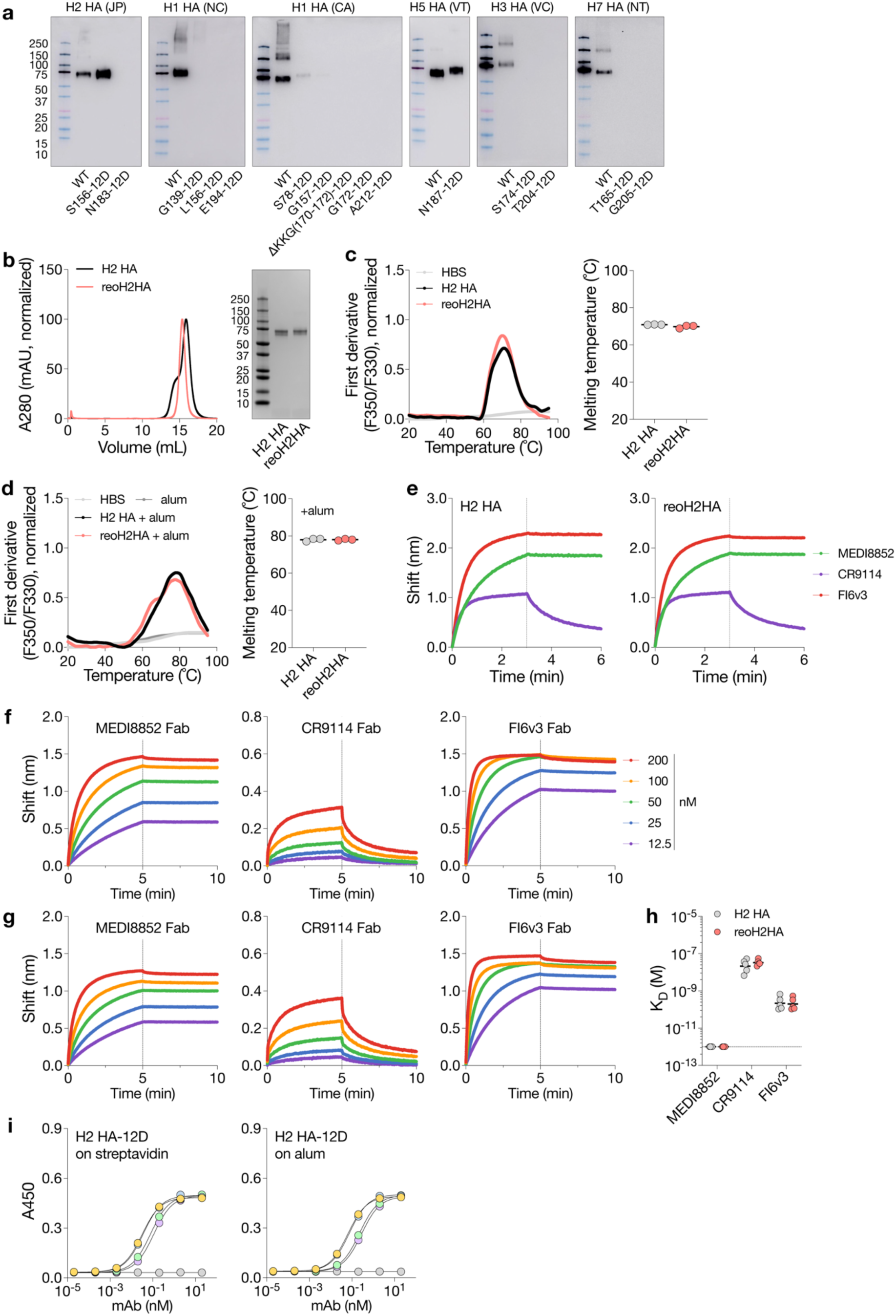
OligoD insertion into HA-head of H2 HA. **a**, Screening of oligoD insertion to the head region of different HAs by Western-blot analysis. Blots were detected by a mouse anti-His Tag antibody. HA sequences were from H1 HA (NC), H1 HA (CA), H2 HA (JP), H5 HA (VT), H3 HA (VC – A/Victoria/3/1975) or H7 HA (NT – A/FPV/Dutch/1927). **b**, Size-exclusion purification and gel electrophoresis of wild-type H2 HA and reoH2HA. Gels were stained with Coomassie brilliant blue. **c,d**, Thermal melting profiles and T_m_ of wild-type H2 HA and reoH2HA in the absence © or presence of alum (**d**, HA: alum = 1: 10, *w/w*). Center values represent the mean (*n*=3 samples per group). **e**, BLI binding profiles of wild-type H2 HA or reoH2HA with three HA-stem-directed mAbs (MEDI8852, CR9114 and FI6v3). **F,g**, BLI binding profiles of wild-type H2 HA (**f**) or reoH2HA (**g**) with the fragment antigen-binding (Fab) of MEDI8852, CR9114 or FI6v3. Dashed lines indicate the beginning of dissociation in **e-g**. **h**, Binding affinities (K_D_) of stem-directed antibodies to wild-type H2 HA or reoH2HA calculated based on data in **f,g**. Data are presented as the geometric mean of the log-transformed K_D_ values. Dashed line indicates the limit of quantification. **i**, Binding of head-directed (8F8 and 8M2) or stem-directed (MEDI8852 and FI6v3) mAbs to H2 HA-12D (H2 HA with 12D inserted into its C-terminus) on streptavidin-(left) or alum-based (right) ELISA plates. Data are presented as mean ± s.d. (*n*=2 biological replicates of technical duplicates).

**Figure S9.**
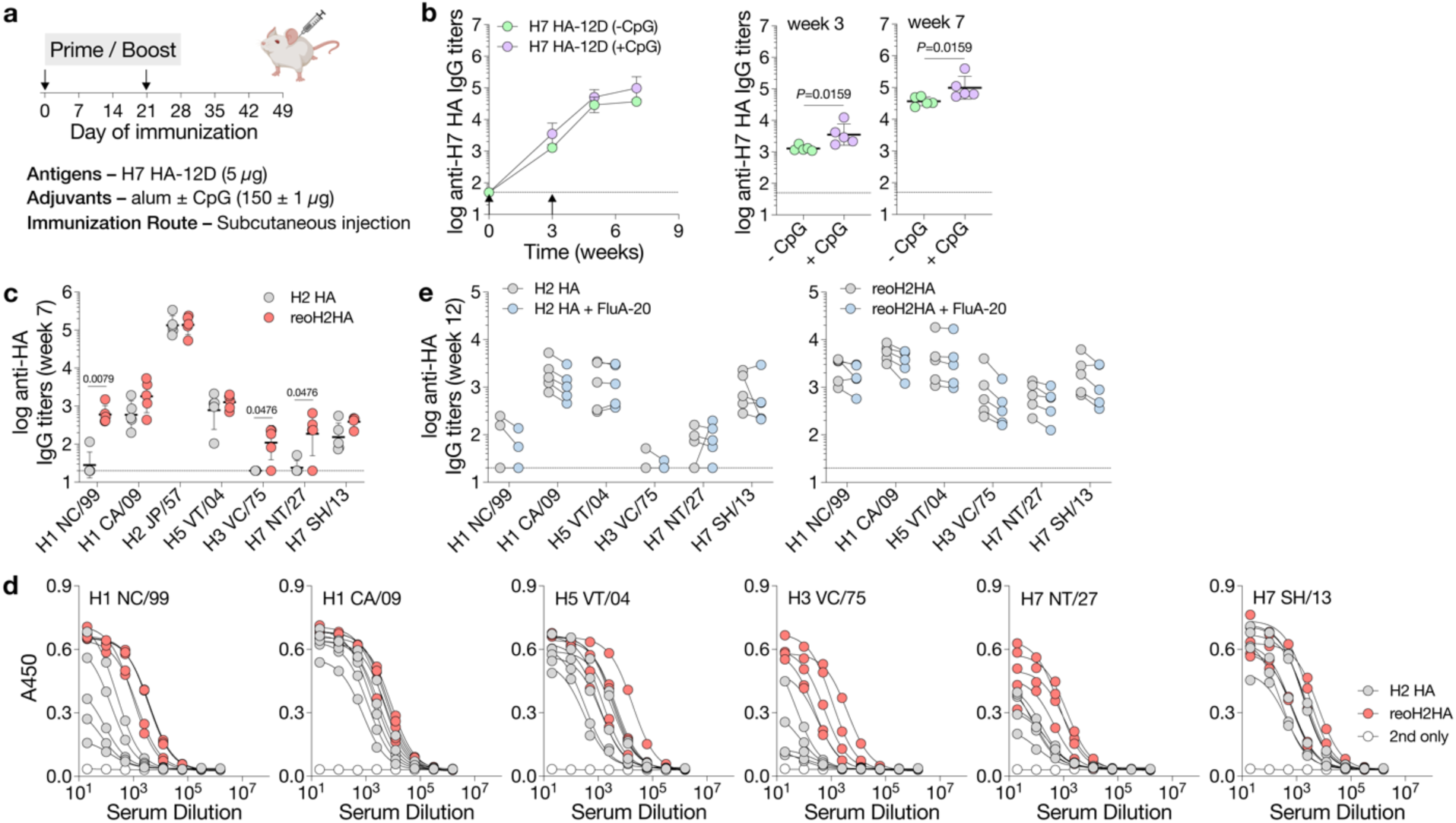
Immunization of oligoD-modified HA. **a**, A prime-boost immunization study with H7 HA-12D adjuvanted with alum alone or alum/CpG *via* subcutaneous injection in BALB/c mice (*n*=5 mice per group). **b**, Serum H7 HA-specific IgG titers over time. Antibody titers of week 3 and 7 from the two groups are plotted on the right for comparison. Data are presented as the geometric mean ± s.d. of the log-transformed values. **c**, Cross-reactive binding of different group 1 (H1 NC/99, H1 CA/09, H2 JP/57, H5 VT/04) and group 2 HAs (H3 VC/75, H7 NT/27, H7 SH/13 – A/Shanghai/2/2013) by week-7 antisera (*n*=5 mice per group). Data are presented as the geometric mean ± s.d. of the log-transformed values. **d**, ELISA binding curves of week-12 antisera against different group 1 and 2 HAs. **e**, Serum binding titers to different group 1 (H1 NC/99, H1 CA/09, H2 JP/57, H5 VT/04) and group 2 HAs (H3 VC/75, H7 NT/27, H7 SH/13) in the presence of a competing mAb – FluA-20. Data are presented as the geometric mean of the log-transformed values (*n*=5 mice per group). Dashed lines indicate the limit of quantification in **b,c,e**. Comparison of two groups was performed using the two-tailed Mann–Whitney U test. *P* values of 0.05 or less were considered significant and plotted.

